# Illuminating the Role of A-to-I Editing in Cancer using EndoVIA 2.0

**DOI:** 10.1101/2025.09.29.679362

**Authors:** Alexandria L. Quillin, Tatiana F. Flores, Devanshi C. Purohit, Angela Halstead, Benoît Arnould, Aastha, Eleonora Grandi, Mason Leffler, José B. Sáenz, Jennifer M. Heemstra

## Abstract

Adenosine-to-inosine (A-to-I) RNA editing is an essential post-transcriptional modification catalyzed by adenosine deaminases acting on RNA (ADAR) enzymes, and emerging evidence suggests its dysregulation can contribute to cancer. However, technical barriers have hindered spatial analysis of editing activity in formalin-fixed paraffin-embedded (FFPE) tissues – an abundant but challenging sample type. Here, we introduce EndoVIA 2.0, an optimized immunostaining assay that enables quantification and spatial mapping of edited RNAs in FFPE tissues. Using human Endonuclease V (hEndoV) as a molecular affinity reagent, we developed a calcium-dependent staining workflow compatible with crosslinked tissues. EndoVIA 2.0 was validated in cell lines having altered ADAR1 expression, fresh frozen (FF) tissues, gastric organoids, and FFPE tissue sections. Using quantitative imaging combined with machine-learning segmentation and spatial analysis, EndoVIA 2.0 successfully detected A-to-I editing changes in ADAR1 knockout (KO) and overexpression models, revealing differential editing patterns across tissues. In gastric organoids and FFPE tissues, the assay also spatially resolved editing heterogeneity and distinguished ADAR1-deficient from ADAR1-sufficient cells. Application to long-archived lung and breast cancer FFPE tissues suggest EndoVIA 2.0’s broad utility and potential for capturing disease-associated changes in A-to-I editing. Finally, staining across the gastric neoplastic cascade revealed stage-dependent increases in A-to-I editing and changes in subcellular localization, further supporting the possibility that ADAR plays a critical role in the early development of gastric cancer. EndoVIA 2.0 enables robust, spatial detection of A-to-I editing in FFPE tissues – circumventing the limitations of RNA extraction and unlocking access to archived clinical specimens. This platform lays the foundation for mapping RNA editing dynamics in cancer progression and may support future biomarker discovery.

## Introduction

Adenosine-to-inosine (A-to-I) RNA editing is among the most abundant and essential post-transcriptional modifications in eukaryotes.[1] Catalyzed by adenosine deaminases acting on RNA (ADARs) within double-stranded regions of RNA (dsRNA), A-to-I editing plays a critical role in modulating the innate immune response.[1] Of the three mammalian ADARs, ADAR1 is ubiquitously expressed and serves as the primary editor. ADAR2, while present in other tissues, is most active in the brain and ADAR3 is catalytically inactive.[2] As dysregulated A-to-I editing continues to emerge as a crucial factor in tumorigenesis, the opportunity to probe inosine in clinically relevant tissues will aid in the detection, diagnosis, and management of human cancer.

From a clinicopathologic perspective, formalin-fixed paraffin-embedded (FFPE) tissues are invaluable for cancer diagnosis and represent the vast majority of samples in long-term pathology archives.[3] A-to-I editing is routinely characterized using RNA sequencing (RNA-seq) because reverse transcriptases recognize inosine as guanosine.[4] However, formalin fixation causes RNA-protein crosslinking and degradation that complicate RNA extraction, often resulting in degraded RNA that is unfit for acquiring robust sequencing data.[5,6] Sequencing fragmented, low-quality RNA increases the likelihood of sequence misalignments, random sampling errors, and false positive reads that incorrectly identify unedited sites as edited.[7,8] Furthermore, extraction-based methods erase cell-to-cell variation, which is essential for identifying tumor heterogeneity.[9] Although *in situ* approaches could overcome these limitations, reliable anti-inosine antibodies for direct detection of edited RNA in FFPE tissues are unavailable.[10–12] Consequently, clinically relevant archived specimens remain inaccessible using current A-to-I editing characterization techniques.

To address this challenge, we have harnessed a conserved nucleic acid repair enzyme, Endonuclease V (EndoV), that recognizes and cleaves inosine-containing transcripts in the presence of magnesium.[13] In contrast, calcium supports inosine binding without cleavage, enabling EndoV to function as an inosine affinity reagent.[14,15] Most recently, we developed Endonuclease V Immunostaining Assay (EndoVIA), which leverages EndoV for *in situ* fluorescent detection and quantification of A-to-I editing while preserving the subcellular localization of inosine-containing transcripts and providing insight into tissue heterogeneity.[16] We therefore sought to adapt EndoVIA to fresh frozen (FF) and FFPE tissues to enable the characterization of A-to-I editing in countless archived samples.

Multiple studies have demonstrated abnormal expression of ADAR enzymes and dysregulated A-to-I editing in various cancers, including gastric cancer (GC).[17–21] Recent work by Sáenz and colleagues has provided additional insight into the role of ADAR in gastric tumorigenesis.[22,23] Their studies revealed dsRNA accumulation during gastric epithelial injury and metaplastic reprogramming, implicating ADAR1-mediated A-to-I editing in the regulation of these responses.[22,23] These findings highlight the need to examine the spatial distribution of A-to-I editing across gastric disease progression, positioning GC as a relevant setting to demonstrate the capabilities of EndoVIA 2.0.[24]

In this work, we present EndoVIA 2.0, an optimized immunostaining workflow for detecting A-to-I editing in FFPE tissues (Figure 1A). We perform optimization and validation experiments in cell lines and FF tissue and demonstrate that EndoVIA 2.0 is compatible with gastric organoids and clinically archived FFPE samples across a variety of human cancers. Together, these results establish EndoVIA 2.0 as a robust approach for spatially mapping A-to-I editing in clinically relevant tissues, expanding our ability to investigate the landscape of RNA editing across human health and disease.

**Figure 1.**
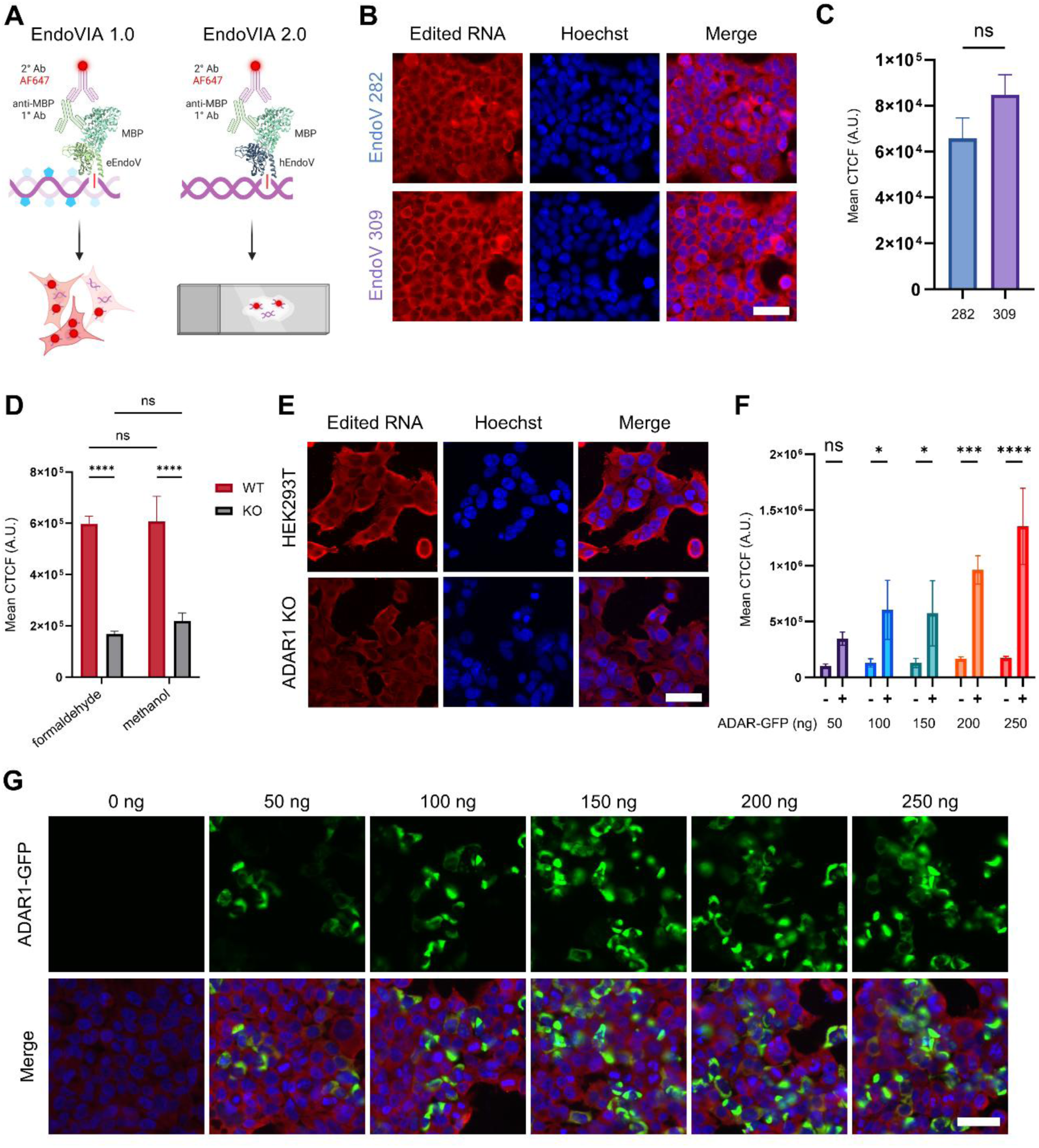
Leveraging human EndoV to detect A-to-I editing *in situ.* (A) Schematic of EndoVIA using *E. coli* EndoV (eEndoV) to detect inosine-containing transcripts in cells, and EndoVIA 2.0 using human EndoV (hEndoV) for detection in crosslinked FFPE tissues. (B) Fixed HEK293T cells were immunostained with hEndoV 282 or hEndoV 309 for edited RNA and cell nuclei (Hoechst; blue). Scale bar, 50 μm. (C) Quantification of mean corrected total cellular fluorescence (CTCF) from (B). (D) Mean CTCF of edited RNA in WT and ADAR1 KO HEK293T cells fixed with either formaldehyde or methanol. (E) WT and ADAR1 KO HEK293T cells stained with EndoVIA 2.0 and Hoechst (cell nuclei; blue). Scale bar, 50 μm. (F) Quantification of mean CTCF of edited RNA in ADAR1-GFP positive (+) and negative (-) cells from (G). (G) HEK293T cells transfected with increasing amounts of GFP-tagged ADAR1-p110 plasmid (0–250 ng) and then stained to detect edited RNA (red) using EndoVIA 2.0 and cell nuclei (Hoechst; blue). Scale bar, 50 μm. Data are representative of three independent experiments. Data are shown as mean ± s.d. in arbitrary units (A.U.). Statistical analysis: unpaired *t*-test (C), one-way ANOVA (D), two-way ANOVA (F). ns, not significant, \**P* < 0.0332, \*\*\**P* < 0.0002, \*\*\*\**P* < 0.0001. Created in BioRender. Quillin, A. (2026) https://BioRender.com/85ysbws

## Materials and Methods

### Ethical Approval and Patient Consent

All mouse experiments were performed according to protocols approved by the Washington University in St. Louis School of Medicine Animal Studies Committee. Human gastric specimens were obtained under protocols approved by the Institutional Review Boards at the Universidad del Cauca and Washington University in St. Louis School of Medicine (IRB 201901176), or from the Digestive Diseases Research Core Center Biobank (IRB 201111078). Written informed consent was obtained as approved by the appropriate Institutional Review Board. Additional human tissues were obtained from the Siteman Cancer Center Tissue Procurement Core under an approved biospecimen use agreement. Samples were provided coded and de-identified, and investigators had no access to participant identifiers. The Washington University Human Research Protection Office determined that this study did not constitute human subjects research (IRB 202303104).

### EndoVIA 2.0

EndoVIA 2.0 was performed on cultured cells, murine gastroids, FF tissues, and FFPE tissues as described below, with detailed protocols provided in the Supplementary Materials and Methods. Samples were fixed, permeabilized, and blocked prior to incubation with MBP-tagged hEndoV-282 in calcium-containing blocking buffer at 150 nM. Samples were washed, incubated with anti-MBP primary antibody, followed by Alexa Fluor 647-conjugated secondary antibody and Hoechst nuclear stain. Importantly, calcium was maintained in all buffers during and following hEndoV incubation.

### Microscopy and Image Analysis

Fluorescence imaging was performed using a Nikon Spinning-Disk microscope (Nikon Instruments Inc., Melville, NY, USA). For each experiment, acquisition settings were established using the sample with the highest fluorescence signal to avoid saturation and were held constant across all samples within the experimental set. Images were analyzed using Cellpose 2.0 and Fiji (ImageJ, 1.54 p) to quantify mean fluorescence intensity or corrected total cellular fluorescence (CTCF).[26]

Gastroid images were segmented using Cellpose 2.0. A pretrained TissueNet model was retrained on EndoVIA 2.0-stained gastroid images using EndoVIA and Hoechst fluorescence to generate single-cell segmentation masks. Masks were used to extract EndoVIA and Ai9 fluorescence intensity and centroid position for each segmented cell. Cells were classified into populations based on EndoVIA fluorescence, and classifications were mapped back to the corresponding cellular positions. Additional details of the segmentation and analysis workflow are provided in the Supplementary Materials and Methods.

For tissue analyses, cellular regions of interest (ROIs) were generated by segmentation of the E-cadherin channel. EndoVIA and E-cadherin fluorescence intensities were measured within each ROI and background-subtracted. Background-corrected ROI measurements were averaged within each tissue section, and normalized EndoVIA fluorescence was calculated as the mean background-corrected EndoVIA fluorescence divided by the corresponding mean background-corrected E-cadherin fluorescence. Section-level values were averaged to obtain one patient-level value for each independent case. CTCF was additionally calculated as integrated density – (ROI area x mean background fluorescence).

### Statistics

Statistical analyses were completed using GraphPad Prism 10 (GraphPad Software, Boston, MA, USA) with the statistical tests specified in the corresponding figure legends.

## Results

### Development of EndoVIA 2.0

In our initial workflow (EndoVIA 1.0), inosine-containing RNA was detected in fixed cells using *Escherichia coli* EndoV (eEndoV) fused to a maltose-binding protein (MBP) tag, followed by staining with primary anti-MBP and fluorescent secondary antibodies. While effective in cultured cells, we recognized that significant modifications to the protocol would be required to enable robust detection in FFPE tissues. Because ADAR-mediated editing occurs within structured duplexes, inosines are most often found in dsRNA. We have observed that eEndoV has reduced affinity for these substrates and prefers inosine-containing single-stranded RNA (ssRNA). We initially addressed this by incorporating a denaturation step with glyoxal, which disrupts Watson-Crick-Franklin base pairing and leads to unfolding. While this approach proved successful in the development of EndoVIPER, EndoVLISA, and EndoVIA 1.0, we recognized that it would be less suited for extending EndoVIA into tissue samples, given that FFPE tissues are extensively crosslinked from formalin fixation and that RNA unfolding may not be feasible.

EndoV is a highly conserved enzyme, present in all forms of life.[28] We therefore hypothesized that other homologs may have increased affinity and selectivity for inosine in dsRNA, eliminating the need for denaturation. We determined that human EndoV (hEndoV) was particularly promising. Alternative splicing of the *ENDOV* transcript produces multiple variants; however, only a small subset encodes full-length proteins containing the conserved catalytic core required for inosine recognition. Among these, the canonical isoform (hEndoV 282) and two larger isoforms (hEndoV 308 and hEndoV 309) represent catalytically active variants reported in human cells.[29] We cloned, purified, and expressed both the hEndoV 282 and hEndoV 309 isoforms with the same MBP tag that was used in EndoVIA 1.0. We fixed HEK293T cells with formaldehyde and performed EndoV immunostaining with both constructs and quantified the fluorescence. Importantly, staining was performed in the presence of calcium to promote binding to the inosine-containing RNA. We determined that both isoforms yielded detectable signal with minimal background (Figure 1B) and no significant difference in quantified fluorescence (Figure 1C). As a result, we chose to move forward with hEndoV 282 for subsequent experiments.

Adenosine deaminases acting on tRNA (ADAT) independently modify adenosines within the anticodon loop, making tRNA one of the most abundant sources of inosine-containing ssRNA.[30] Because eEndoV has preference for inosine within ssRNA, there was concern during the development of the original EndoVIA workflow that the presence of tRNA could interfere with our ability to detect ADAR-mediated editing events occurring within dsRNA structures. To address this in EndoVIA 1.0, we incorporated methanol fixation to reduce tRNA retention and glyoxal denaturation to disrupt RNA secondary structure, providing eEndoV access to inosines within dsRNA. Using tRNA fluorescence *in situ* hybridization (FISH), we previously demonstrated that methanol fixation and washing reduced detectable tRNA by ∼72% relative to formaldehyde fixation across two cell lines.[16] In our experiments, hEndoV has been shown to recognize inosine-containing RNA in double-stranded contexts. As additional validation, we compared EndoVIA 2.0 staining in HEK293T cells fixed with either formaldehyde or methanol. Staining revealed comparable fluorescence across both conditions, providing strong evidence that the presence of tRNA does not interfere with hEndoV-mediated detection of ADAR-edited transcripts (Figure 1D).

We next validated the specificity of EndoVIA 2.0 for edited RNA by altering ADAR1 levels in cells. We observed a significant decrease in edited RNA fluorescence in ADAR1 knockout (ADAR1 KO) cells relative to wild-type (WT) HEK293T cells (Figure 1E). WT cells exhibited ∼5-fold higher fluorescence than ADAR1 KO cells, providing greater discrimination than previously observed using EndoVIA 1.0. Importantly, the global editing levels of these WT and ADAR1 KO cell lines were previously characterized by performing RNA sequencing, which demonstrated a decrease in Alu Editing Index (AEI) values from 1.4 in WT cells to 0.2 in ADAR1 KO cells.[16] Conversely, overexpression of green fluorescent protein (GFP)-tagged ADAR1-p110, the constitutively expressed ADAR1 isoform, led to significantly higher fluorescence in transfected cells compared to non-transfected cells (Figures 1F and 1G, supplementary material, Figure S1). To quantify the magnitude of fluorescence differences detected by EndoVIA 2.0 following ADAR1 overexpression, we calculated the ratio of mean EndoVIA fluorescence in transfected cells relative to non-transfected cells. Across the tested conditions, transfected cells exhibited approximately 3.4- to 7.7-fold higher EndoVIA fluorescence than non-transfected cells (supplementary material, Figure S2). Additionally, we used EndoVIA 2.0 to detect A-to-I editing in a malignant cell line that has been reported to display hypo-editing (supplementary material, Figure S3). Consistent with our previous observations using EndoVIA 1.0, EndoVIA 2.0 remained compatible with super-resolution imaging as well. dSTORM imaging of HEK293T cells revealed increased fluorescence near the cell membrane, similar to the localization pattern previously observed using EndoVIA 1.0, with minimal signal detected in the absence of hEndoV (supplementary material, Figure S4).[16] Together, these results demonstrate that hEndoV can be used to specifically detect and quantify edited RNA in formaldehyde-fixed cells without the need for RNA denaturation, laying the foundation for its application to FFPE tissues.

### EndoVIA 2.0 in Tissues

We adapted EndoVIA 2.0 for FFPE compatibility by first applying the workflow to FF tissues, which preserves RNA integrity and allows optimization in high-quality samples. To ensure our workflow modification to enable EndoV binding did not interfere with immunostaining performance in tissues, we stained two protein controls – β-actin and histone H4 – and observed strong fluorescence intensity and appropriate subcellular localization (supplementary material, Figure S5). We then assessed editing across tissue types with reported differences in ADAR activity. We selected brain and adrenal gland as representative models of high and low editing levels.[31] EndoVIA 2.0 immunostaining and quantification revealed significantly higher edited RNA fluorescence in brain tissue (Figures 2A and 2B).

**Figure 2.**
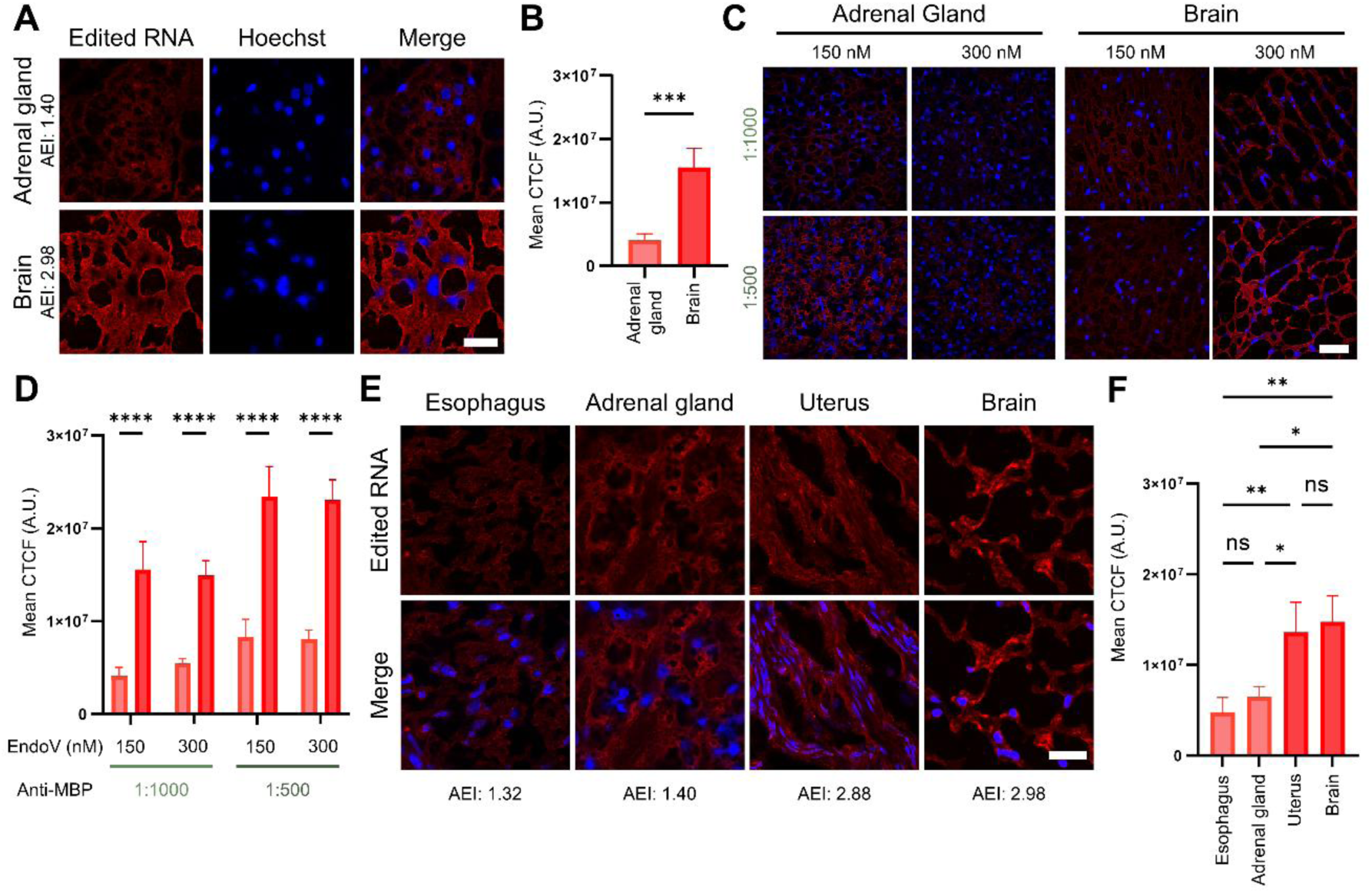
Staining edited RNA in FF tissues with varying levels of A-to-I editing. (A) Human adrenal gland and brain FF tissues were immunostained using EndoVIA 2.0 and Hoechst (blue; nuclei). Scale bar, 25 μm. (B) Quantification of mean CTCF from (A). (C) Human adrenal gland and brain FF tissues were stained with hEndoV (150 or 300 nM) and anti-MBP antibody (1:1000 or 1:500). Hoechst (blue) was used to visualize nuclei. Scale bar, 50 μm. (D) Quantification of mean CTCF from (C). hEndoV concentrations are indicated below each condition, and anti-MBP antibody dilutions are indicated in green. (E) Human esophagus, adrenal gland, uterus, and brain FF tissues were immunostained using EndoVIA 2.0 and Hoechst (blue; nuclei). Scale bar, 25 μm. (F) Quantification of mean CTCF from (E). Data are representative of three independent experiments. Data are shown as mean ± s.d. in arbitrary units (A.U.). Statistical analysis: unpaired *t*-test (B), one-way ANOVA (D, F). ns, not significant, \**P* < 0.0322, \*\**P* < 0.0021, \*\*\**P* < 0.0002, \*\*\*\**P* < 0.0001.

To optimize staining conditions, we next evaluated a range of antibody concentrations across both high- and low-editing tissues (Figure 2C). MBP was used in place of hEndoV as a negative control to assess background signal (supplementary material, Figure S6). While increasing the concentration of hEndoV did not improve staining, higher concentrations of anti-MBP primary antibody enhanced fluorescence intensity (supplementary material, Figure S7). Importantly, all tested conditions yielded staining patterns consistent with known differences in editing levels between tissues (Figure 2D). Based on these results, we selected 150 nM hEndoV and 1:500 (2 μg/mL) anti-MBP for subsequent experiments. To further evaluate EndoVIA 2.0 across tissues with differing levels of A-to-I editing, we examined esophagus and uterus alongside adrenal gland and brain. Consistent with previously reported differences in global A-to-I editing activity, EndoVIA 2.0 fluorescence was higher in brain and uterus compared with adrenal gland and esophagus (Figures 2E and 2F). Notably, the reported AEI values are approximately 1.32 and 1.40 for esophagus and adrenal gland, and 2.88 and 2.98 for uterus and brain, respectively.[31]

### EndoVIA 2.0 in Gastric Samples

To expand our workflow to the *in vitro* setting, we leveraged an organoid model system. Organoids offer a physiologically relevant, three-dimensional model that recapitulates key features of native tissue architecture and cellular heterogeneity, making them an ideal system for probing spatial patterns of RNA editing in a biologically meaningful context. We sought to apply EndoVIA 2.0 to validated gastric organoid models. We relied on gastric organoids (gastroids) isolated from *Adar1^fl/fl^;Mavs^-/-^;Ai9* mice, where *Adar1* can be inducibly deleted following transduction with an adenoviral *Cre* vector. The *Mavs^-/-^* background was used to suppress the downstream MAVS-dependent response to dsRNA accumulation and permit viable *Adar1-*deficient gastric epithelial cells to be studied.[23] In our *in vitro* system, *Adar1^fl/fl^;Mavs^-/-^;Ai9* gastroids were transduced with a Cre recombinase-expressing adenoviral vector to inducibly delete *Adar1*. Importantly, Cre-mediated deletion of *Adar1* is marked by the expression of the Ai9 fluorescent reporter, enabling identification of cells within the heterogeneous organoid population that have undergone *Adar1* deletion. Following staining with EndoVIA 2.0 and subsequent imaging, we observed lower edited RNA fluorescence in Ai9-positive (*Adar1*-deficient) cells compared to Ai9-negative (*Adar1*-sufficient) cells, consistent with a loss of A-to-I editing activity (Figure 3A, supplementary material, Figure S8).

**Figure 3.**
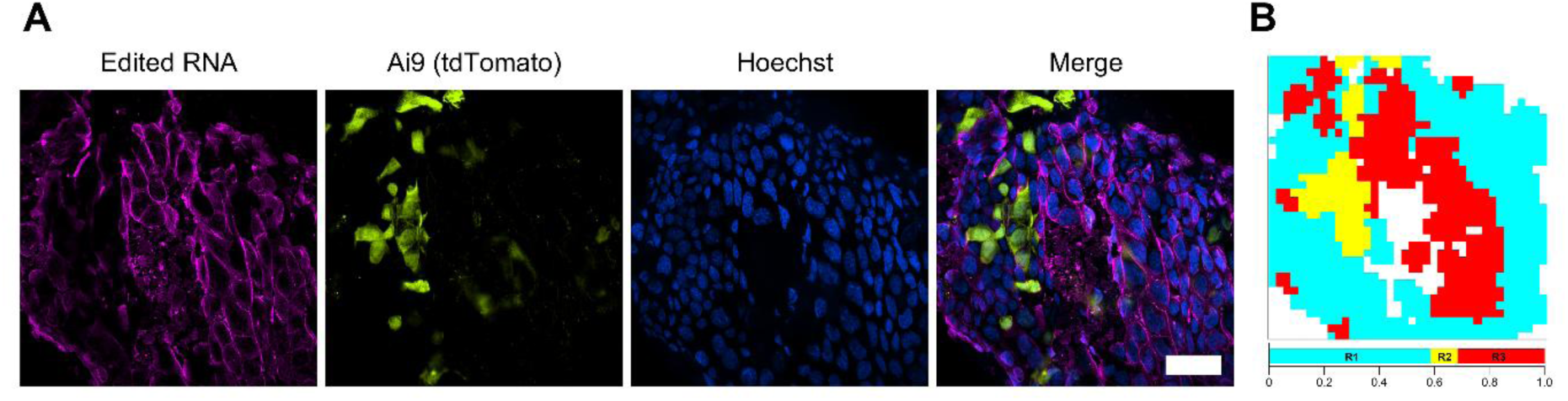
Spatial quantification of A-to-I editing in murine gastric organoids. (A) *Adar1^fl/fl^;Mavs^-/-^;Ai9* gastroids were transduced with a Cre recombinase-expressing adenoviral vector to induce *Adar1* deletion in Ai9-positive cells. Organoids were immunostained using EndoVIA 2.0 to detect edited RNA, *Adar1*-deficient parietal cells (Ai9), and cell nuclei (blue). Scale bar, 50 μm. (B) Cells were phenotyped using predefined intensity thresholds in the EndoVIA channel as hypo-editing (R2, yellow), hyper-editing (R3, red), or intermediate/undefined (R1, blue; cells not meeting the criteria for R2 or R3). Data are representative of three independent experiments.

To quantify editing at the single-cell level, we utilized computational image analysis to perform segmentation and extract mean fluorescence intensity values for both Ai9 and EndoVIA channels for each cell. These measurements were plotted for individual cells to visualize the relationship between Cre-mediated recombination and edited RNA signal (supplementary material, Figure S9). Single-cell phenotyping was then performed using EndoVIA fluorescence to classify cells into three distinct populations that were generated based on predefined intensity thresholds: hypo-editing (R2), hyper-editing (R3), and an intermediate/undefined group that did not meet the predefined criteria for either group (R1). These classifications were subsequently mapped back to the spatial location of each gastroid (Figure 3B). Importantly, the hypo-editing population (R2) was defined independently of Ai9 status and corresponded with the *Adar1*-deficient cells, confirming the specificity of EndoVIA 2.0 for detecting A-to-I editing. R1 represents cells with intermediate EndoVIA signal and therefore was not expected to be uniformly positive or negative for edited RNA. Together, these results highlight the ability of EndoVIA 2.0 to reveal spatial patterns of RNA editing heterogeneity in complex tissue models and demonstrate its compatibility with single-cell analysis workflows. This approach lays the groundwork for spatial phenotyping and molecular stratification based on RNA editing signatures.

To validate EndoVIA 2.0 in FFPE tissues with known differences in ADAR1 activity, we applied our workflow to gastric corpus sections from *Adar1^ΔPC^;Mavs^-/-^* mice, which lack ADAR1 expression in acid-secreting parietal cells. Similarly to our approach used for gastroids, the *Mavs*^-/-^ background was used to limit MAVS-dependent innate immune activation triggered by ADAR1 loss, thereby allowing the effect of ADAR1 deficiency on RNA editing to be assessed in viable gastric epithelial cells. These mice transgenically express a parietal cell-specific *Cre* recombinase, allowing detection of *Adar1*-deficient parietal cells using the *Ai9* reporter. Specifically, detection of the reporter (Ai9, tdTomato) enables internal comparison between *Adar1*-deficient (tdTomato^+^) and *Adar1*-sufficient cells within the same tissue section. Tissues were stained, imaged, and individual cells were segmented to extract the mean fluorescence intensities for tdTomato and edited RNA (Figures 4A and 4B).

**Figure 4.**
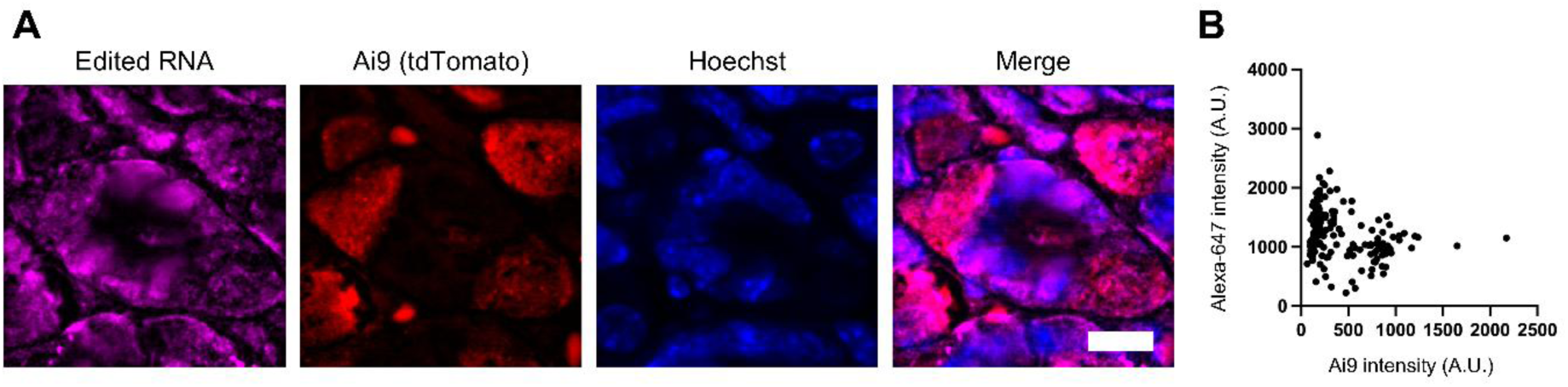
Detecting A-to-I editing in murine *Adar1^ΔPC^;Mavs^-/-^* FFPE tissues. (A) FFPE gastric corpus sections were co-immunostained using EndoVIA 2.0 to detect edited RNA, parietal cell-specific Ai9 reporter, cell nuclei (blue), and cell membranes (green, not shown). Scale bar, 50 μm. (B) Quantification of single-cell fluorescence of edited RNA and Ai9. Data are representative of three biologically independent gastric corpus sections.

As expected, tdTomato-negative (*Adar1*-sufficient) cells exhibited higher edited RNA fluorescence than their tdTomato-positive (*Adar1*-deficient) counterparts. Consistent with these findings, independent RNA sequencing analysis of *Adar1*-sufficient and *Adar1*-deficient parietal cells from the *Adar1^ΔPC^* model demonstrated a substantial loss of ADAR1-dependent editing following *Adar1* deletion.[32] This independent molecular characterization further supports the reduction in EndoVIA 2.0 fluorescence observed in *Adar1*-deficient cells. Notably, some EndoVIA signal was still detected in the tdTomato-positive population. Given the loss of ADAR1-dependent editing shown by RNA sequencing and the residual fluorescence in stained FFPE sections omitting EndoV, we interpret this remaining signal in *Adar1*-deficient cells as likely being immunostaining-associated background commonly observed in immunohistochemistry (Figure S10).[33] EndoVIA 2.0 fluorescence also varied across individual cells within the tdTomato-negative group, suggesting potential heterogeneity of A-to-I editing within gastric epithelium.

### EndoVIA 2.0 Across Human Cancer Types

To evaluate the broader applicability of EndoVIA 2.0 beyond gastric models, we stained matched malignant and non-malignant FFPE tissue samples from participants with lung cancer or breast cancer – two cancer types previously associated with hyper-editing.[19] Given that these specimens had been archived for over 20 years, we first assessed tissue integrity by immunostaining cytoplasmic (β-actin) and nuclear (histone H4) markers. Both markers yielded sufficient signal and appropriate localization, supporting preserved tissue architecture and antigenicity (supplementary material, Figure S11). Interestingly, fluorescence from these control markers was higher in malignant tissues than matched non-malignant tissue. To account for potential differences in overall staining intensity between specimens, we therefore incorporated E-cadherin, an epithelial membrane-associated protein, as an internal reference in the EndoVIA 2.0 workflow.

EndoVIA fluorescence was detectable in both malignant and non-malignant breast and lung tissues (Figures 5A and 5C).

**Figure 5.**
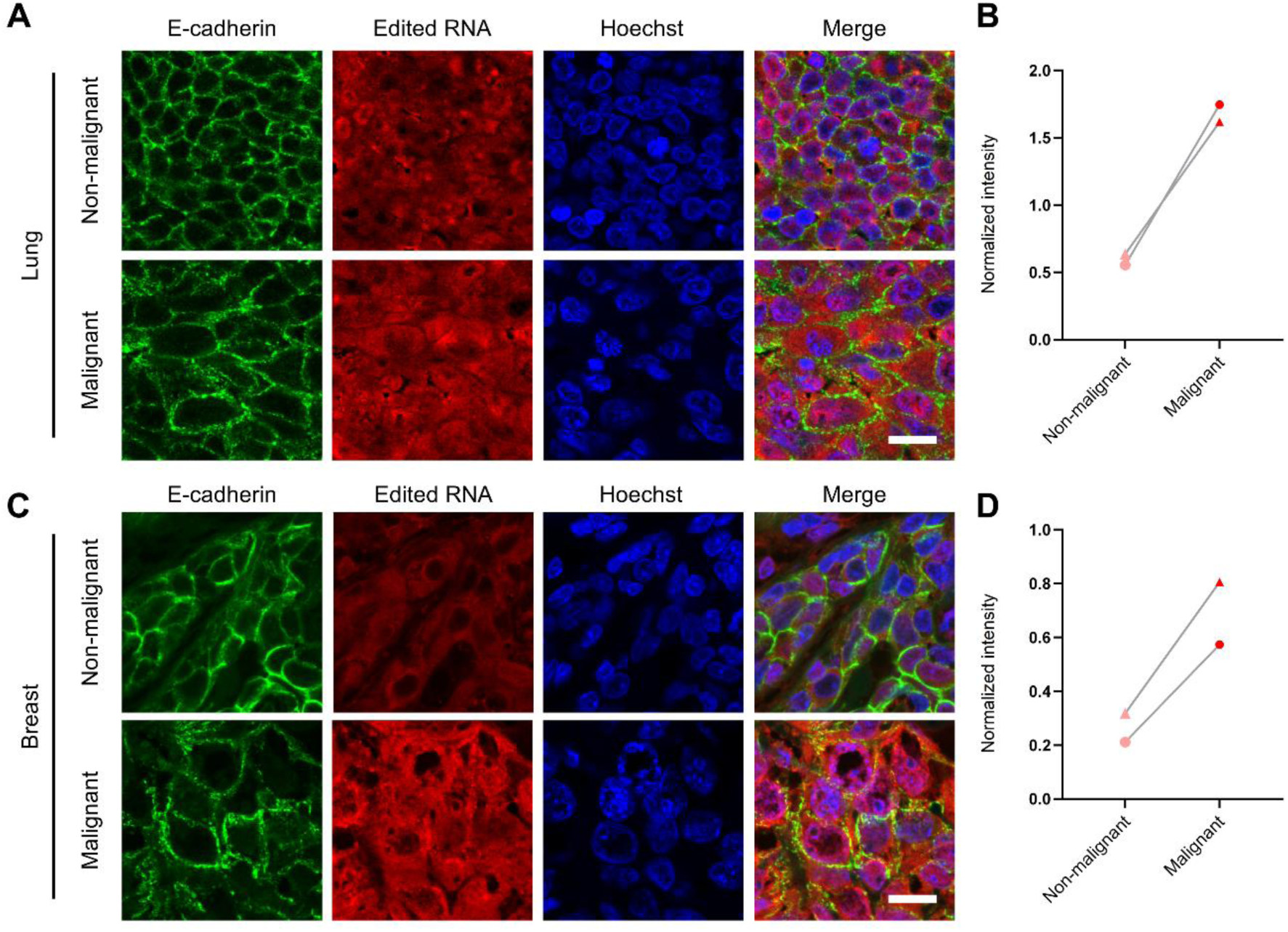
Staining edited RNA in malignant and non-malignant FFPE tissues. (A) Human malignant and non-malignant lung FFPE tissues immunostained for edited RNA using EndoVIA 2.0, E-cadherin, and cell nuclei (Hoechst). Scale bars, 50 μm. (B) Quantification of mean normalized edited RNA intensity from (A). (C) Human malignant and non-malignant breast FFPE tissues immunostained for edited RNA using EndoVIA 2.0, E-cadherin, and cell nuclei (Hoechst). Scale bars, 50 μm. (D) Quantification of mean normalized edited RNA intensity from (C). Each point represents the patient-level mean from one independent case, and lines connect matched non-malignant and malignant tissues from the same patient. Two independent matched patient cases were analyzed for each tissue type (breast, n = 2; lung, n = 2). Multiple tissue sections, acquired images, and cellular regions of interest contributed to each patient-level value. Data are shown as mean ± s.d.

Normalized EndoVIA fluorescence was higher in malignant tissue relative to matched non-malignant tissue in both breast cases and lung cases (Figure 5B and 5D). To determine whether E-cadherin normalization influenced the observed differences, we evaluated EndoVIA and E-cadherin fluorescence separately. E-cadherin fluorescence exhibited modest case-dependent differences between matched non-malignant and malignant tissues. In contrast, EndoVIA fluorescence was higher in malignant tissue in both breast cases and both lung cases prior to normalization, consistent with previously reported increases in A-to-I editing in these cancer types.[19,34,35] As an additional analysis independent of E-cadherin normalization, we quantified corrected total cellular fluorescence (CTCF) for EndoVIA. EndoVIA CTCF was similarly higher in malignant relative to matched non-malignant tissue in both breast cases and both lung cases, indicating that the observed difference was not dependent on E-cadherin normalization or a single fluorescence quantification approach (Figure 5, supplementary material, Figures S12 and S13). Together, these preliminary results demonstrate that EndoVIA 2.0 is compatible with diverse FFPE tissue types, including long-archived clinical samples, and can detect disease-associated differences in EndoVIA fluorescence across multiple cancer types.

### Mapping A-to-I Editing Across the Gastric Neoplastic Cascade

Having determined that EndoVIA 2.0 enables detection of edited RNA in human FFPE tissues across multiple cancer types, we next explored whether our approach could be used to characterize the changing landscape of A-to-I editing during gastric disease progression. Chronic inflammation is a well-established precursor to gastric tumorigenesis, and recent studies from the Sáenz lab have shown that dsRNA accumulates in inflamed and pre-neoplastic gastric epithelium. These findings suggest that A-to-I editing may play an important role in modulating dsRNA levels during early stages of disease. However, the spatial distribution and extent of A-to-I editing across these stages remain poorly understood. To further examine the interplay between A- to-I editing and gastric disease progression, we used EndoVIA 2.0 to stain human gastric FFPE tissues spanning the gastric neoplastic cascade, including normal tissue, chronic non-atrophic gastritis (CNG), chronic atrophic gastritis (CAG), CAG with intestinal metaplasia (CAG/IM), and gastric adenocarcinoma (Figure 6A). Three independent patient cases were analyzed for each histopathological group.

**Figure 6.**
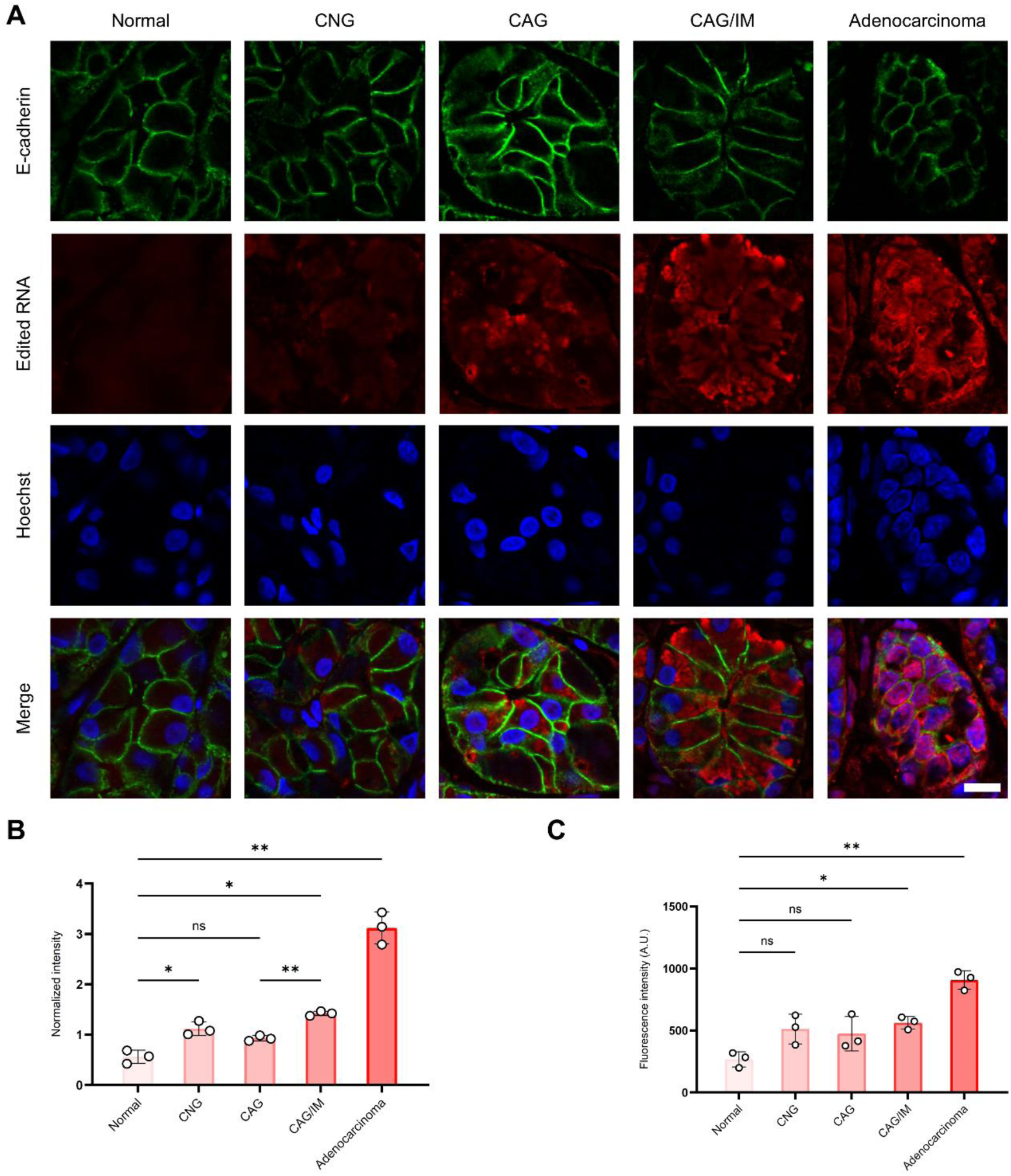
Staining edited RNA across the gastric neoplastic cascade. (A) Human FFPE gastric tissues representing normal gastric epithelium, chronic non-atrophic gastritis (CNG), chronic atrophic gastritis (CAG), CAG with intestinal metaplasia (CAG/IM), and gastric adenocarcinoma (cancer) immunostained for edited RNA using EndoVIA 2.0, E-cadherin, and cell nuclei (Hoechst). Scale bar, 10 μm. (B) Quantification of mean normalized edited RNA intensity from (A) across normal, CNG, CAG, CAG/IM, and cancer tissues. (C) Quantification of mean EndoVIA fluorescence across the same histopathological groups prior to E-cadherin normalization. Three independent patient cases were analyzed for each histopathological group (n = 3 patients/group; 15 patients total). Multiple tissue sections, acquired images, and cellular regions of interest contributed to each patient-level value. Data are shown as mean ± s.d. Statistical analysis was performed using Welch’s one-way ANOVA followed by Dunnett’s T3 multiple-comparisons test. For normalized edited RNA intensity in (B), Welch’s ANOVA demonstrated a significant overall difference among groups (W(4, 4.705) = 57.53, *P* = 0.0003). Significant Dunnett’s T3 adjusted *P* values were normal versus CNG, 0.0414; normal versus CAG/IM, 0.0325; normal versus cancer, 0.0052; CNG versus cancer, 0.0109; CAG versus CAG/IM, 0.0017; CAG versus cancer, 0.0271; and CAG/IM versus cancer, 0.0442. ns, not significant; \**P* < 0.05; \*\**P* < 0.01.

Quantification of normalized EndoVIA 2.0 signal differed significantly across the stages of gastric disease (Figure 6B). Compared to normal tissue, fluorescence was significantly higher in CNG, CAG/IM, and gastric adenocarcinoma, whereas the difference between CAG and normal tissue was not significant. CAG/IM also exhibited significantly higher fluorescence than CAG. Gastric adenocarcinoma exhibited the highest signal and was significantly elevated relative to each of the other histopathological groups. Analysis of EndoVIA and E-cadherin fluorescence further demonstrated that the observed pattern was not solely attributable to E-cadherin normalization (Figure 6C, and supplementary material, Figure S14).

Notably, in tissues corresponding to intermediate stages of disease, including CNG, CAG, and CAG/IM, EndoVIA signal frequently appeared as bright, subcellular structures that were largely absent in normal gastric epithelium and appeared more diffuse in gastric adenocarcinoma samples. Although the identity and biological significance of these structures remain unclear, their distinct spatial distribution suggests that further characterization of the subcellular localization of edited RNA across gastric disease progression may be warranted. Together, these findings demonstrate that EndoVIA 2.0 enables spatial detection of A-to-I editing across stages of gastric disease and reveal differences in EndoVIA fluorescence and spatial distribution among the histopathological groups examined.

## Discussion

In this work, we developed EndoVIA 2.0, a next-generation immunostaining workflow designed to detect and quantify inosine-containing transcripts in crosslinked cells and tissues. By leveraging hEndoV, we established a robust workflow capable of spatially resolving A-to-I editing events in FFPE tissues. Through systematic validation in cell lines, organoids, and both FF and FFPE tissues, we demonstrated that EndoVIA 2.0 is sensitive to changes in ADAR1 expression, compatible with 3D and archival tissue models, and capable of revealing cellular heterogeneity in editing levels. Together, these results establish the specificity, flexibility, and spatial resolution of EndoVIA 2.0 across diverse biological contexts.

EndoVIA 2.0 enables *in situ* detection of A-to-I editing in FFPE tissues, overcoming a longstanding limitation of existing methods. Current approaches for characterizing editing events typically require RNA extraction, which is often unreliable in FFPE samples due to degradation, crosslinking, and loss of spatial context. In contrast, our approach preserves tissue architecture and enables relative quantification of editing at the single-cell level within intact tissue sections. EndoVIA 2.0 complements quantitative and site-specific characterization provided by sequencing-based approaches, enabling a more comprehensive view of A-to-I editing. This not only expands the experimental landscape for studying A-to-I editing but also unlocks access to an extensive archive of well-annotated, patient-derived samples that were previously inaccessible using traditional methods.

The ability to visualize and quantify A-to-I editing in FFPE tissues also opens new opportunities to investigate the relationship between RNA editing and disease progression. Using both engineered organoid models and gastric tissue, we demonstrated that EndoVIA 2.0 can detect editing differences within heterogeneous epithelial populations and distinguish between ADAR1-deficient and ADAR1-sufficient cells. Consistent with emerging evidence linking ADAR activity to gastric tumorigenesis, EndoVIA fluorescence differed across stages of the gastric neoplastic cascade, with the highest signal observed in gastric adenocarcinoma. Interestingly, inflammatory stages exhibited distinct subcellular structures suggesting accumulation of edited RNA that were not prominent in normal tissue. These findings demonstrate the ability of EndoVIA 2.0 to visualize differences in both abundance and spatial distribution of inosine-containing RNA across gastric disease states and provide a framework for future studies investigating the biological significance of these patterns. Although these clinical analyses demonstrate the applicability of EndoVIA 2.0 to archived human FFPE tissues, the patient cohorts examined herein remain modest. Validation in larger, independent cohorts will therefore be necessary to determine the reproducibility and generalizability of the observed disease-associated differences.

Ultimately, this workflow illuminates a new dimension of insight into the landscape of A-to-I editing in human disease. Given the challenges of detecting cancer at early stages and the growing interest in RNA-based biomarkers, the ability to assess editing directly *in situ* within FFPE tissues holds strong potential for identifying new indicators of disease risk, progression, or therapeutic response. While additional biological studies will be needed to further define these relationships, these results establish EndoVIA 2.0 as a powerful platform for spatial detection of A-to-I editing in clinically relevant contexts and lay the foundation for future studies aimed at deciphering the role of RNA editing in cancer biology, biomarker discovery, and therapeutic development.

## Supporting information

EndoVIA 2.0 SI

## Acknowledgements

This work was supported by the National Institutes of Health (R35GM144075 to J.M.H.), National Institute of Diabetes and Digestive and Kidney Diseases (NIDDK; R01141682 to J.B.S), Digestive Diseases Research Core Center (Washington University in St. Louis School of Medicine), Tissue Procurement Core (Washington University in St. Louis School of Medicine), and the Gastric Cancer Research Innovation Fund (Foundation for Barnes-Jewish Hospital). The content is solely the responsibility of the authors and does not necessarily reflect the official views of the listed supporters. Illustrations were created with BioRender.com. We would also like to thank Dr. Charles Rice for gifting us the ADAR1 KO cell line and Kevin Kaifer for guidance with image acquisition.

## Author Contributions Statement

ALQ, JBS, and JMH conceived and designed the study. ALQ, TFF, JBS, and JMH contributed to the development of the EndoVIA 2.0 workflow. ALQ, TFF, DCP, and A carried out experiments. BA developed and conducted specialized image analysis. A performed super-resolution imaging. DCP, EG, and ML expressed and purified human EndoV used in the assay. AH and JBS provided gastric samples and contributed to biological insight and interpretation of data. ALQ, TFF, and BA analyzed the data. ALQ, TFF, BA, and A generated figures. ALQ and TFF wrote the original draft of the manuscript. All authors contributed to the interpretation of the data, reviewed and edited the manuscript, and approved the final version of the manuscript. JBS and JMH supervised the study and secured funding.

## Data Availability Statement

The data that support the findings of this study are available from the corresponding authors upon reasonable request.

## Abbreviations

ADARs: adenosine deaminases acting on RNA
ADATs: adenosine deaminases acting on tRNA
AEI: Alu Editing Index
A-to-I editing: adenosine-to-inosine editing
A.U.: arbitrary units
CAG: chronic atrophic gastritis
CAG/IM: CAG with intestinal metaplasia
CNG: chronic non-atrophic gastritis
CTCF: corrected total cellular fluorescence
dSTORM: direct stochastic optical reconstruction microscopy
dsRNA: double-stranded RNA
EndoV: Endonuclease V
EndoVIA: Endonuclease V Immunostaining Assay
eEndoV: *Escherichia coli* EndoV
FF: fresh frozen
FISH: fluorescence *in situ* hybridization
FFPE: formalin-fixed paraffin-embedded
GC: gastric cancer
GFP: green fluorescent protein
hEndoV: human Endonuclease V
KO: knockout
MBP: maltose-binding protein
RNA-seq: RNA sequencing
ROI: region of interest
ssRNA: single-stranded RNA
TIRF: total internal reflection fluorescence microscopy
WT: wild type

## Supplementary Material Online

**EndoVIA 2.0_Journal of Pathology_SI**

**Supplementary Materials and Methods**

### Supplementary Figures

**Figure S1.** Detection of edited RNA in cells transfected with ADAR1-GFP.

**Figure S2.** Fold difference in EndoVIA 2.0 fluorescence between ADAR1-GFP-positive and ADAR1-GFP-negative cells.

**Figure S3.** Detection of hypo-editing in malignant kidney cells.

**Figure S4.** Super-resolution microscopy of edited RNA using EndoVIA 2.0.

**Figure S5.** Detection of β-actin and Histone H4 in fresh frozen uterus tissue.

**Figure S6.** Assessing non-specific binding of antibody concentrations.

**Figure S7.** Determination of optimal EndoV and anti-MBP concentrations.

**Figure S8.** Staining edited RNA in murine gastric organoids.

**Figure S9.** Mean fluorescence intensities of edited RNA and Ai9.

**Figure S10.** No-hEndoV control for FFPE EndoVIA 2.0 staining.

**Figure S11.** Detection of β-actin and Histone H4 in lung FFPE tissues.

**Figure S12.** Analysis of E-cadherin and EndoVIA 2.0 fluorescence in lung tissues.

**Figure S13.** Analysis of E-cadherin and EndoVIA 2.0 fluorescence in breast tissues.

**Figure S14.** Analysis of E-cadherin fluorescence across the gastric neoplastic cascade.

### Supplementary Tables

**Table S1.** Buffer composition of various solutions used in Methods.

**Table S2.** Source of antibodies and dyes used in Methods.

