## Supplementary material for "Illuminating the Role of A-to-I Editing in Cancer using EndoVIA 2.0": EndoVIA 2.0 SI

#### Supplementary Materials

#### Table of Contents

| | |
|---|---|
| **Supplementary Materials and Methods** | **S2-S5** |
| **Supplementary Figures** | |
| **Figure S1.** Detection of edited RNA in cells transfected with ADAR1-GFP. | **S6** |
| **Figure S2.** Fold difference in EndoVIA 2.0 fluorescence between ADAR1-GFP-positive and ADAR1-GFP-negative cells. | **S7** |
| **Figure S3.** Detection of hypo-editing in malignant kidney cells. | **S8** |
| **Figure S4.** Super-resolution microscopy of edited RNA using EndoVIA 2.0. | **S9** |
| **Figure S5.** Detection of β-actin and Histone H4 in fresh frozen uterus tissue. | **S10** |
| **Figure S6.** Assessing non-specific binding of antibody concentrations. | **S11** |
| **Figure S7.** Determination of optimal EndoV and anti-MBP concentrations. | **S12** |
| **Figure S8.** Staining edited RNA in murine gastric organoids. | **S13** |
| **Figure S9.** Mean fluorescence intensities of edited RNA and Ai9. | **S14** |
| **Figure S10.** No-hEndoV control for FFPE EndoVIA 2.0 staining. | **S15** |
| **Figure S11.** Detection of β-actin and Histone H4 in lung FFPE tissues. | **S16** |
| **Figure S12.** Analysis of E-cadherin and EndoVIA 2.0 fluorescence in breast tissues. | **S17** |
| **Figure S13.** Analysis of E-cadherin and EndoVIA 2.0 fluorescence in breast tissues. | **S18** |
| **Figure S14.** Analysis of E-cadherin fluorescence across the gastric neoplastic cascade. | **S19** |
| **Supplementary Tables** | |
| **Table S1.** Buffer composition of various solutions used in Methods. | **S20** |
| **Table S2.** Antibodies, dyes, and key reagents used in Methods. | **S21** |

#### Supplementary Methods

Unless otherwise noted, all experiments were performed under nuclease-free conditions, and solutions were prepared fresh for each experiment. All incubations for tissue staining experiments were performed in a humidified chamber.

##### Ethical Approval and Patient Consent

*Mouse Husbandry and Gastric Tissue Collection:* *ROSA26$^{LSLTdTomato}$* (*Ai9* reporter; strain 007909) and *Mavs$^{-/-}$* (strain 008634) mice were obtained from the Jackson Laboratory (Bar Harbor, ME, USA). *Adar1*-floxed mice[25] were provided by Qingde Wang (University of Pittsburgh Medical Center, Pittsburgh, PA, USA). All experiments involving mice were performed according to protocols approved by the Washington University in St. Louis School of Medicine Animal Studies Committee.

Human gastric specimens were obtained either through a collaboration between JBS and the Universidad del Cauca (Popayán, Colombia), with approval from the Institutional Review Boards at the Universidad del Cauca and at the Washington University in St. Louis School of Medicine (IRB 201901176), or from the Digestive Diseases Research Core Center Biobank (IRB 201111078). Written informed consent was obtained as approved by the appropriate Institutional Review Board. Additional human tissue specimens were obtained from the Siteman Cancer Center Tissue Procurement Core at Washington University in St. Louis under an approved biospecimen use agreement. Samples were provided coded and de-identified, and investigators had no access to participant identifiers. The Washington University Human Research Protection Office determined that this study did not constitute human subjects research (IRB 202303104).

Mice were maintained in a specified pathogen-free barrier facility under a 12-hour light cycle and had access to food and water *ad libitum*. Prior to obtaining gastric tissue, mice were euthanized using isoflurane for anesthesia, followed by cervical dislocation.

*Adenoviral Transduction of Murine Gastric Organoids:* Gastric organoids were isolated and cultured from the gastric corpora of *Adar1$^{fl/fl}$*; *Mavs$^{-/-}$;Ai9* mice, aged 6-10 weeks, as previously described.[31] Organoids were adenovirally transduced with Ad-Cre to induce Cre-mediated *Adar1* deletion and Ai9 (tdTomato) reporter expression.[31] The *Mavs$^{-/-}$* background permits analysis of viable *Adar1*-deficient gastric cells while preventing the downstream MAVS-dependent response to accumulated dsRNA.

##### Sample Preparation and EndoVIA 2.0 Immunostaining

*Cells*: G-402 (ATCC, Manassas, VA, USA) and HEK293T (ATCC) cells were cultured according to ATTC recommendations. HEK293T ADAR1 KO were provided by Dr. Charles Rice and cultured under the same conditions as parental HEK293T cells. For cellular immunofluorescence experiments, cells were seeded into poly-D-lysine-coated black 96-well plates and cultured for 48 h after seeding or transfection with an ADAR1-p110-GFP plasmid (Addgene, Watertown, MA, USA) prior to staining.

Cells were fixed with either 4% paraformaldehyde for 15 min or ice-cold 100% methanol for 20 min at 4 °C prior to staining. Cells were washed with 1X PBS, permeabilized for 20 min, washed again with 1X PBS, and blocked for 1 h. Cells were then incubated with

MBP-tagged hEndoV-282 at 150 nM in calcium-containing blocking buffer for 1 h at RT. Following three washes with calcium-containing wash buffer, cells were incubated with anti-MBP primary antibody (1:500, 2 μg/mL) in calcium-containing blocking buffer for 1 h. Cells were washed and subsequently incubated with Alexa Fluor 647-conjugated secondary antibody (1:1000) and Hoechst nuclear stain in calcium-containing blocking buffer for 1 h. Cells were washed with calcium-containing wash buffer prior to imaging. Buffer compositions and staining reagents are provided in supplementary material, Tables S1 and S2.

*Gastroids*: Gastroids were cultured in Cultrex basement membrane extract as previously described. Once organoids reached sufficient size, plates were chilled on ice for 5 minutes and washed with ice-cold 1X PBS (without $Ca^{2+}$ or $Mg^{2+}$). Cell Recovery Solution was added, and organoids were gently resuspended using a pre-cut pipette tip. The suspension was rocked at 4 °C and then washed in cold 1X PBS. The pellet was then fixed in freshly prepared paraformaldehyde in 1X PBS followed by three 1X PBS washes. Fixed gastroids were transferred to ice for staining.

For EndoVIA 2.0 staining, fixed gastroids were first blocked at room temperature with gentle rocking for 2 h. Gastroids were pelleted and resuspended in 150 nM hEndoV-MBP-282 in calcium-containing blocking buffer and incubated with rocking in the dark overnight at 4 °C. Gastroids were washed with cold calcium-containing wash buffer and then incubated with anti-MBP antibody (1:1000, 1 μg/mL) in calcium-containing blocking buffer with gentle rocking. Following washing, gastroids were stained with Alexa Fluor 647-conjugated secondary antibody (1:1000) and Hoechst nuclear stain in calcium-containing blocking buffer. Gastroids were washed with cold calcium-containing wash buffer, transferred to microscope slides, and mounted with antifade mounting medium (Thermo Fisher Scientific, Waltham, MA, USA). Slides were allowed to dry in the dark before imaging. Buffer compositions and staining reagents are provided in supplementary material, Tables S1 and S2.

*Fresh Frozen (FF) Tissues*: Frozen tissues were equilibrated at −20 °C prior to sectioning, mounted onto positively charged slides, air-dried, and then fixed in formaldehyde. Slides were washed with 1X TBS, permeabilized, and then blocked. Sections were then incubated with 150 nM hEndoV-MBP-282 in calcium-containing blocking buffer for 1 h at RT. Following three washes with calcium-containing wash buffer, sections were incubated with anti-MBP antibody in calcium-containing blocking buffer for 1 h, washed, and subsequently incubated with Alexa Fluor 647-conjugated secondary antibody (1:1000) and Hoechst nuclear stain in calcium-containing blocking buffer. Following staining and washing, sections were mounted with antifade mounting medium, allowed to cure overnight, and imaged the following day.

*Formalin-fixed paraffin-embedded (FFPE) Tissues*: FFPE tissue sections were deparaffinized using HistoClear solution for 6 min and rehydrated through a graded ethanol series (2 x 3 min; 100%, 90%, 70%, 50%) followed by water. Sections were washed with 1X TBS. Antigen retrieval was performed in 10 mM sodium citrate, 0.05% Tween-20, pH 6.0, using a pressure cooker (High Pressure for 3-5 minutes, allow ~30 min for pressure to decrease following cook time). Following antigen retrieval, sections were washed with 1X TBS and blocked for 1-2 h. Sections were incubated with 150 nM

MBP-tagged hEndoV-282 in calcium-containing blocking buffer for 1 hour at RT, washed three times with calcium-containing wash buffer, and incubated with anti-MBP primary antibody [1:500, 2 µg/mL] for 1 h. Following washing, sections were incubated with Alexa Fluor 647-conjugated secondary antibody [1:1000] and Hoechst nuclear stain for 1 hr. Slides were washed and mounted with antifade mounting medium as described for FF tissues. For no-hEndoV controls, FFPE tissue sections were processed using the complete EndoVIA 2.0 staining workflow with recombinant hEndoV omitted; all subsequent staining and imaging conditions were unchanged.

**Fluorescent Immunohistochemistry**

Fresh-frozen (FF) and formalin-fixed paraffin-embedded (FFPE) tissue sections were prepared for immunostaining as described above, including fixation and/or deparaffinization, rehydration, and antigen retrieval as appropriate. Sections were blocked and incubated with the indicated primary antibodies (1 h at RT), followed by the corresponding fluorescent secondary antibodies and nuclear counterstain (1 h at RT). Antibody identities, suppliers, and working dilutions are provided in supplementary material, Table S2. Following staining, sections were washed and mounted with antifade mounting medium as previously described above.

**Microscopy and Image Analysis**

Widefield and confocal fluorescence images were acquired using a Nikon Spinning-Disk microscope (Nikon Instruments Inc., Melville, NY, USA). For each  For each experimental set, acquisition settings were established using the sample exhibiting the highest fluorescence signal to avoid saturation and were subsequently held constant across all samples within that experiment. Images were analyzed using Fiji/ImageJ (version 1.54p, National Institutes of Health, Bethesda, MD, USA) to quantify mean fluorescence intensity, integrated density, or corrected total cellular fluorescence (CTCF), as appropriate. CTCF was calculated as integrated density − (ROI area × mean background fluorescence).

dSTORM imaging was performed on a Nikon n-STORM/TIRF system (Nikon Instruments Inc., Melville, NY, USA) equipped with an Andor iXon 3 EM-CCD camera (Andor Technology, Belfast Northern Ireland, UK). For each field of view, 30,000-40,000 images were taken acquired. The first 5,000 images were excluded, and the remaining stacks were analyzed using the ThunderSTORM imageJ plugin.[27]

**Single-Cell Segmentation and Quantification in Gastroids**

Single-cell segmentation and quantification of gastroid images were performed using a custom image analysis workflow based on Cellpose 2.0. A pre-trained TissueNet model was iteratively retrained on EndoVIA-stained gastroid images using a human-in-the-loop approach. Segmentation errors generated by the pretrained model were manually corrected, and the model was iteratively retrained to optimize single-cell segmentation. Both EndoVIA and Hoechst fluorescence were used to inform segmentation. The refined model was subsequently applied to experimental images to generate single-cell segmentation masks.

A custom Python workflow was used to extract EndoVIA an Ai9 fluorescence intensity and centroid position for each segmented cell. Cells were classified into R1, R2, and R3 populations based on EndoVIA fluorescence intensity; Ai9 fluorescence was not used to define these populations. Cell classifications were subsequently mapped to the centroid positions of the corresponding segmented cells to visualize their distribution within the gastroid. Ai9 fluorescence was then used to visualize their distribution within the gastroid. Ai9 fluorescence was then used independently to evaluate the relationship between the EndoVIA-defined populations and Cre-mediated *Adar1* deletion.

**Statistics**

Statistical analyses were completed using GraphPad Prism 10 (GraphPad Software, Boston, MA, USA) with the statistical tests used for each analysis specified in the corresponding figure legends. For clinical tissue analyses, individual patients were treated as independent biological replicates, with measurements from multiple tissue sections and images compiled to obtain patient-level values prior to statistical analysis. Gastric specimens were analyzed blinded to histopathological classification. Statistical significance was defined as $P < 0.05$.

#### Supplementary Figures

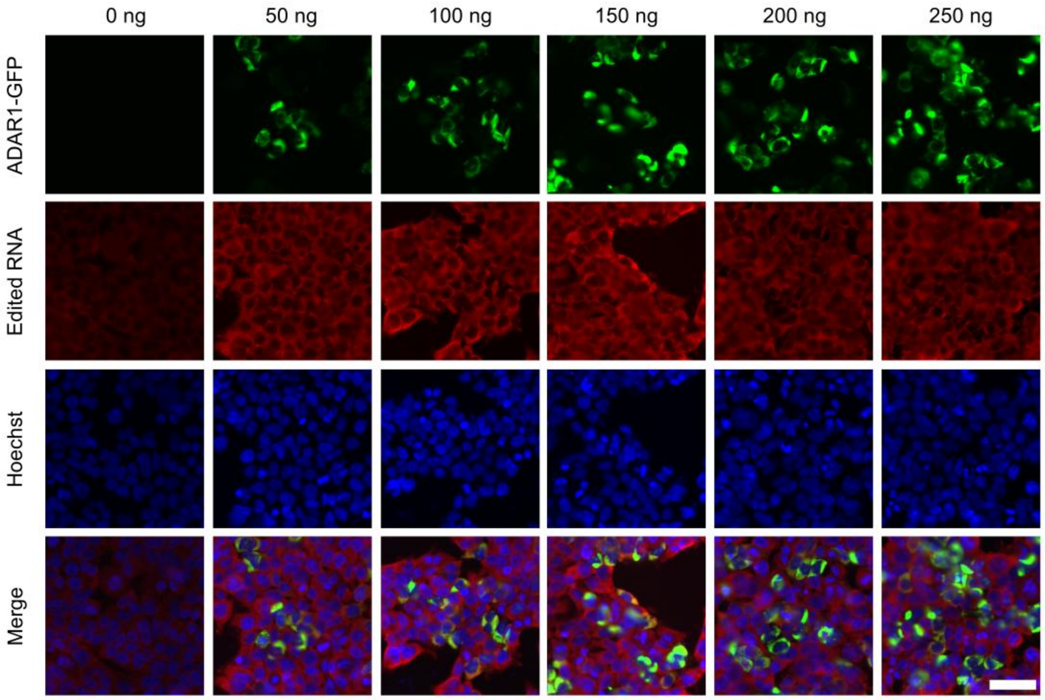


**Figure S1.** Detection of edited RNA in cells transfected with ADAR1-GFP. HEK293T cells transfected with increasing amounts of GFP-tagged ADAR1 p110 plasmid (0–250 ng) and stained for edited RNA using EndoVIA 2.0 and cell nuclei (Hoechst). Scale bar, 50 µm. Data are representative of three independent experiments.

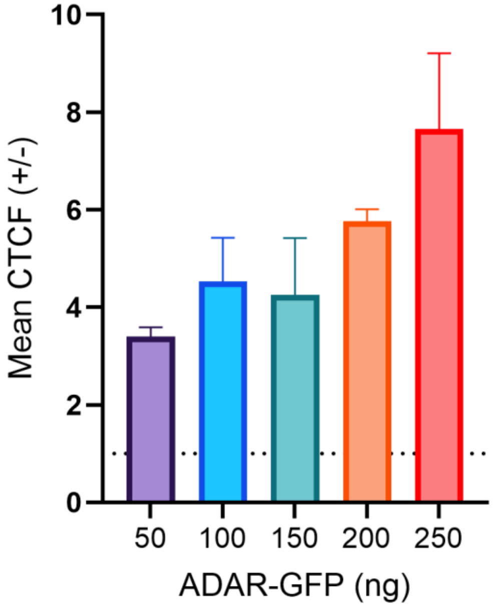


**Figure S2.** Fold difference in EndoVIA 2.0 fluorescence between ADAR1-GFP-positive and ADAR1-GFP-negative cells. HEK293T cells were transfected with increasing amounts of ADAR1-p110-GFP plasmid (50–250 ng), and EndoVIA 2.0 fluorescence was quantified in ADAR1-GFP-positive (+) and ADAR1-GFP-negative (−) cells. For each independent experiment, the mean corrected total cellular fluorescence (CTCF) of ADAR1-GFP-positive cells was divided by the mean CTCF of ADAR1-GFP-negative cells to determine the fold difference in EndoVIA 2.0 fluorescence. Data are representative of three independent experiments. Data are shown as mean ± s.d.

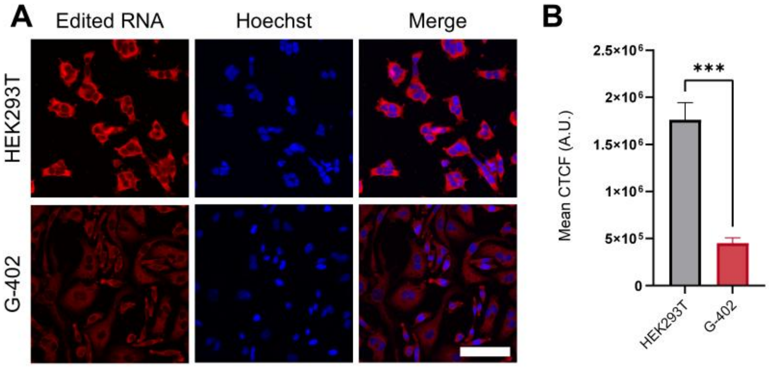


**Figure S3.** Detection of hypo-editing in malignant kidney cells. (A) HEK293T and G-402 cells were immunostained with EndoVIA 2.0 for edited RNA and cell nuclei (Hoechst). Scale bar, 50 μm. (B) Quantification of mean CTCF from (A). Data are representative of three independent experiments. Data are shown as mean ± s.d. in arbitrary units (A.U.). Statistical analysis: unpaired *t*-test. ****$P < 0.0001$.

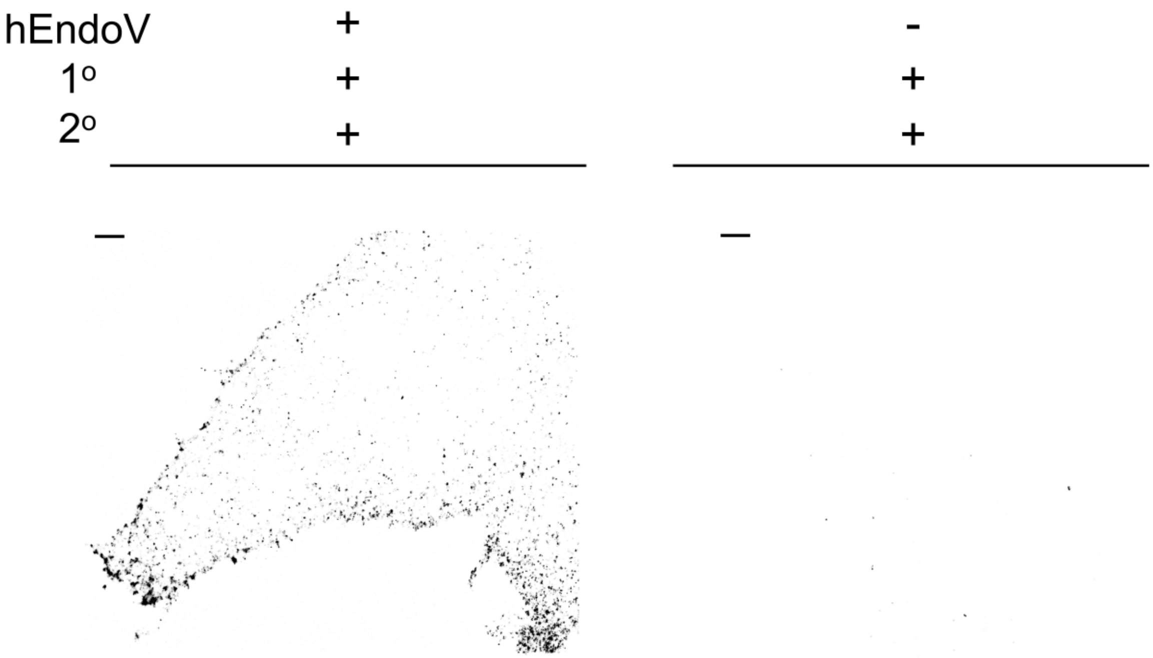


**Figure S4.** Super-resolution microscopy of edited RNA using EndoVIA 2.0. TIRF-dSTORM images of HEK293T cells stained using EndoVIA 2.0 with (+) or without (-) hEndoV. Each image shows a single representative cell. Scale bar, 10 μm. Data are representative of three independent experiments; n=3 coverslips.

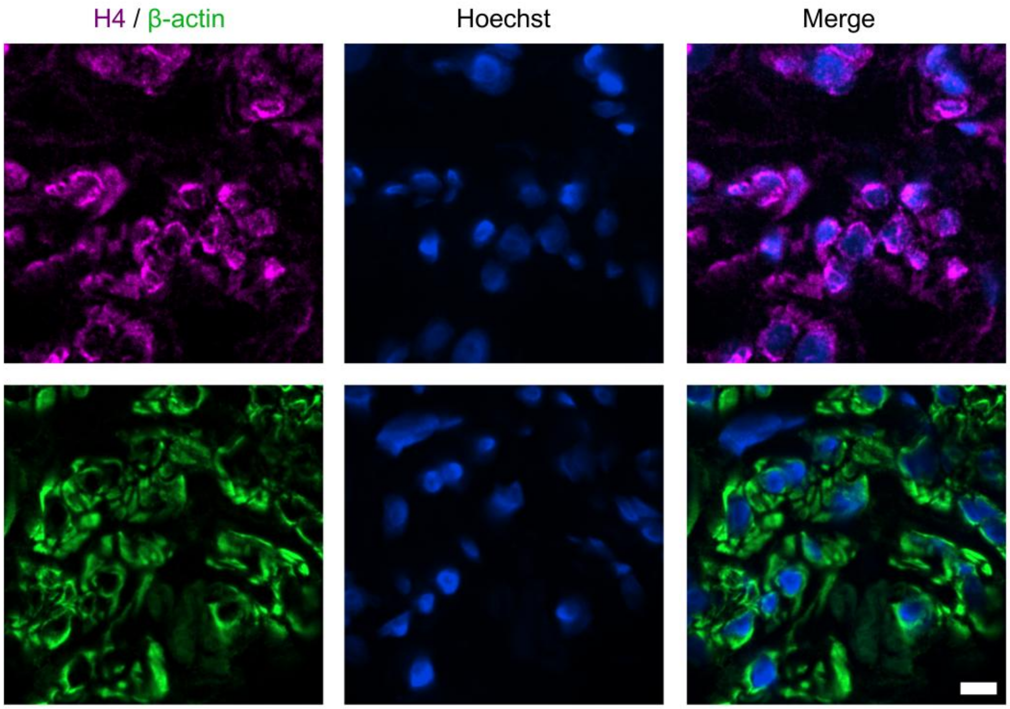


**Figure S5**. Detection of β-actin and Histone H4 in fresh frozen uterus tissue. Human uterus tissue was immunostained for Histone H4, β-actin, and cell nuclei (Hoechst). Scale bar, 5 μm. Data are representative of three independent experiments.

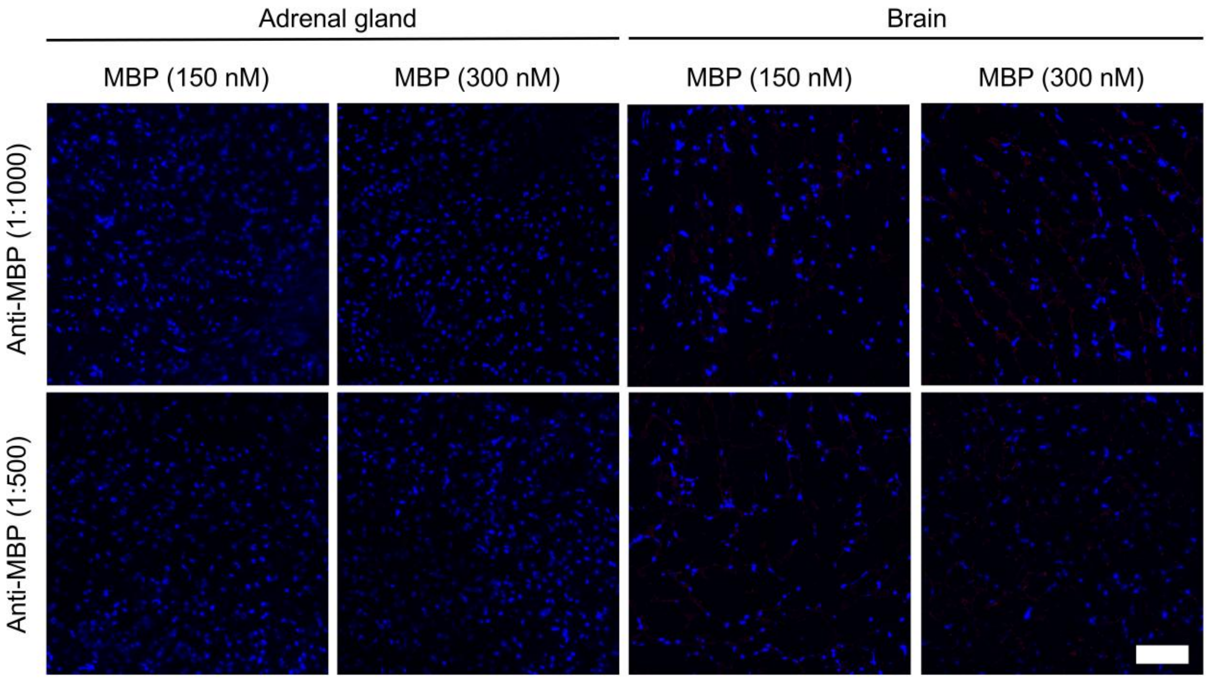


**Figure S6.** Assessing non-specific binding of antibody concentrations. Human adrenal gland and brain fresh frozen tissues were stained with Hoechst (cell nuclei; blue) and increasing amounts of MBP and anti-MBP. Scale bar, 50 µm. Data are representative of three independent experiments.

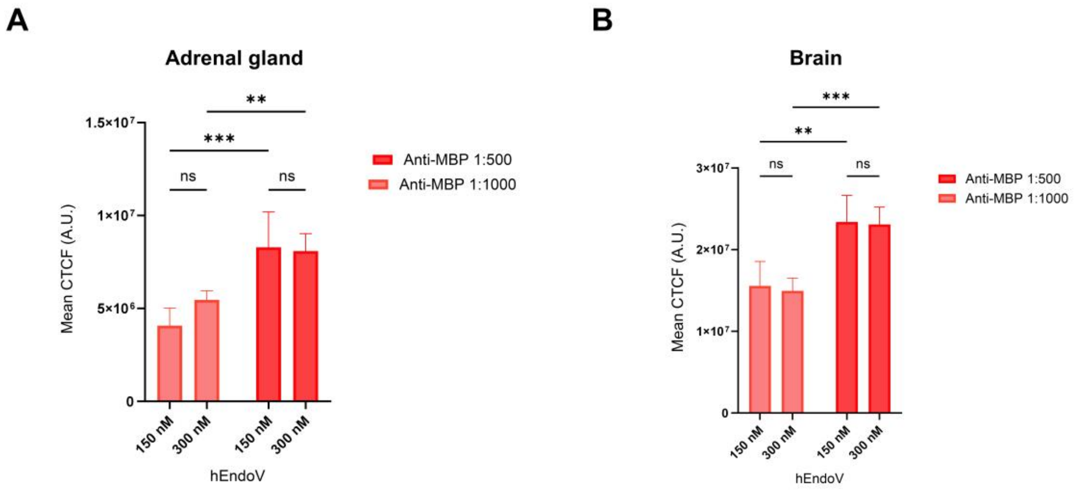


**Figure S7.** Determination of optimal EndoV and anti-MBP concentrations. Human adrenal gland and brain fresh frozen tissues were stained for edited RNA and cell nuclei with increasing amounts of hEndoV or anti-MBP and mean CTCF was quantified. Data are representative of three independent experiments. Data are shown as mean ± s.d. in arbitrary units (A.U.). Statistical analysis: one-way ANOVA (A, B). not significant=ns, ***P* < 0.0021, ****P* < 0.0002.

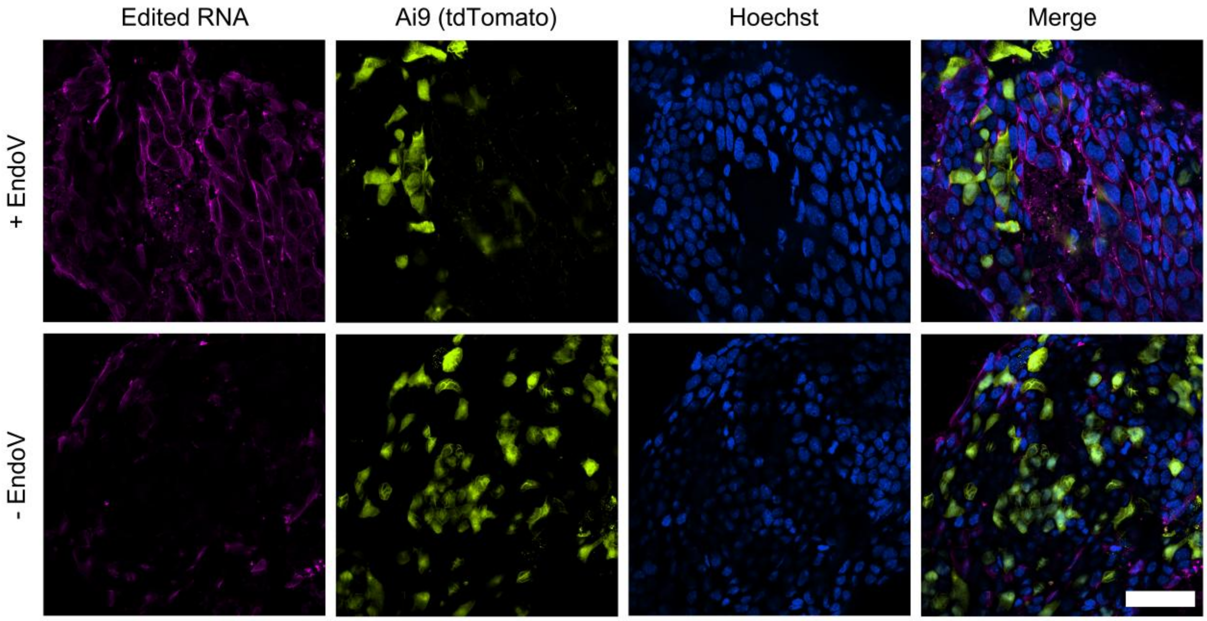


**Figure S8.** Staining edited RNA in murine gastric organoids. *Adar1*$^{fl/fl}$*;Mavs*$^{-/-}$*;Ai9* gastric organoids were transduced with a Cre recombinase-expressing adenoviral vector to induce *Adar1* deletion in Ai9-positive cells. Organoids were immunostained using EndoVIA 2.0 (+EndoV) to detect edited RNA, *Adar1*-deficient parietal cells (Ai9), and cell nuclei (Hoechst). A negative control omitting EndoV (-EndoV) was stained in parallel. Scale bar, 50 µm. Data are representative of three independent experiments.

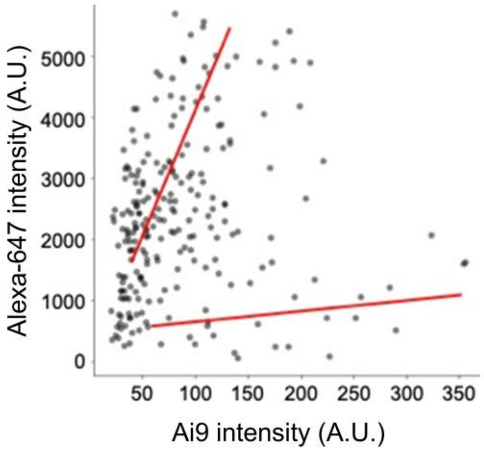


**Figure S9.** Mean fluorescence intensities of edited RNA and Ai9. Single-cell segmentation was performed using a retrained Cellpose 2.0 model, and mean fluorescence intensities were extracted for both edited RNA and Ai9 channels. Data are representative of three independent experiments.

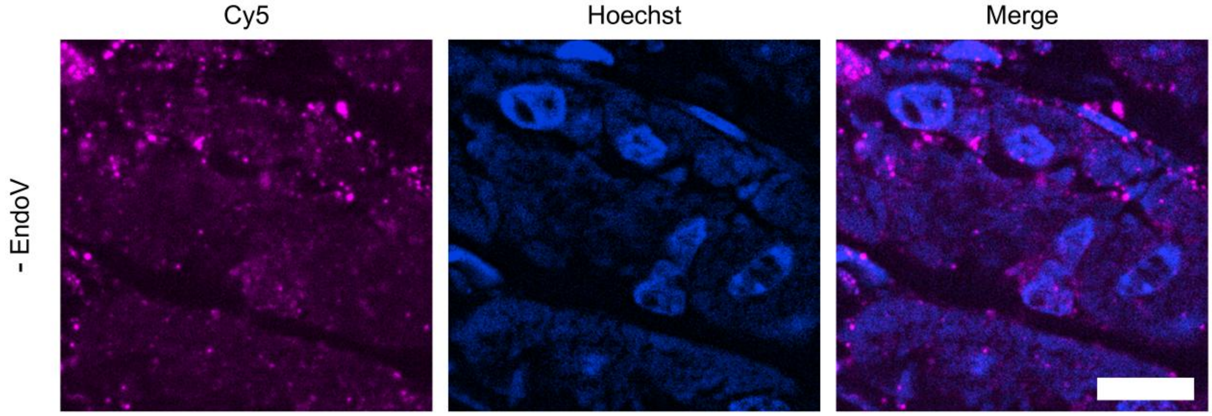


**Figure S10.** No-hEndoV control for FFPE EndoVIA 2.0 staining. Representative FFPE tissue section processed using the complete EndoVIA 2.0 staining workflow with omission of recombinant hEndoV. All other staining reagents and imaging conditions were unchanged relative to EndoVIA-stained samples. Residual fluorescence observed under these conditions represents assay- and tissue-associated background in the absence of the hEndoV recognition reagent. Scale bar, 10 μm. Data are representative of three independent experiments.

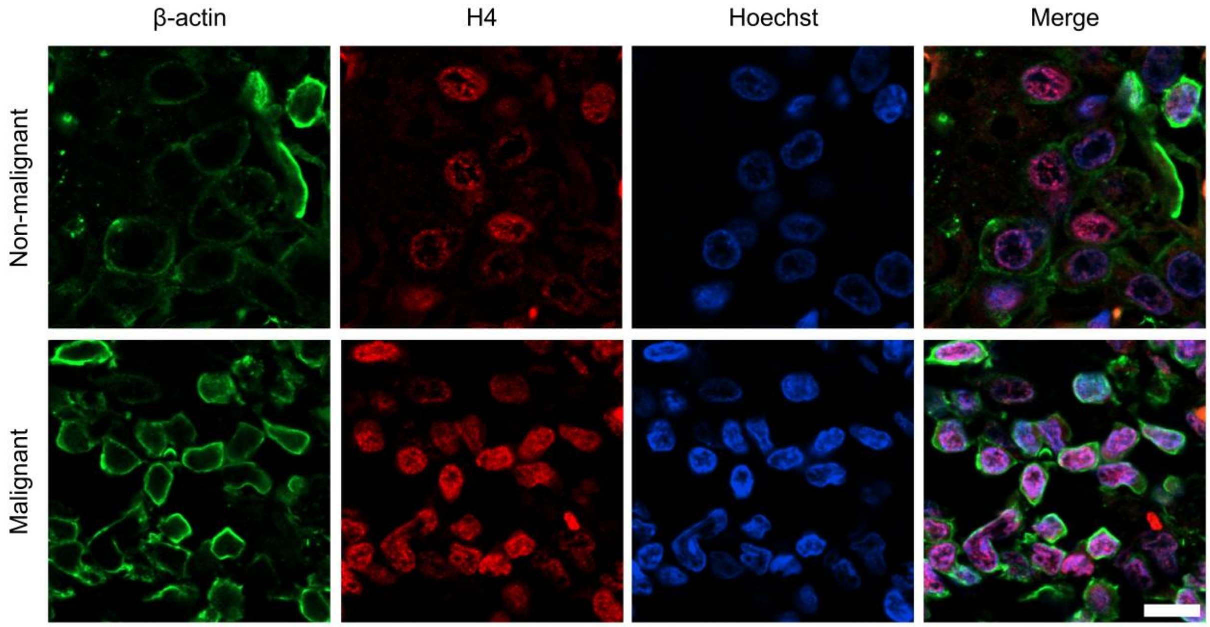


**Figure S11**. Detection of β-actin and Histone H4 in lung FFPE tissues. Human non-malignant and malignant FFPE tissues were immunostained for β-actin, Histone H4, and cell nuclei (Hoechst). Scale bar, 25 μm. Data are representative of three independent experiments.

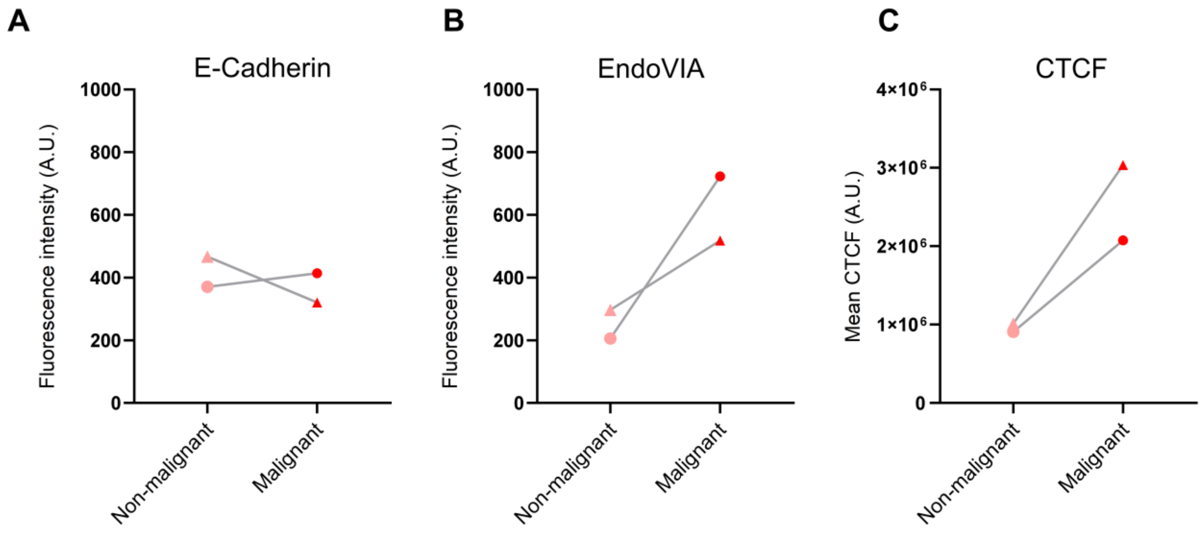


**Figure S12.** Analysis of E-cadherin and EndoVIA 2.0 fluorescence in lung FFPE tissues. (A) Mean E-cadherin fluorescence intensity in matched non-malignant and malignant lung tissues. (B) Mean EndoVIA fluorescence intensity in the same matched tissues. (C) Mean CTCF of EndoVIA in matched non-malignant and malignant tissues. Each point represents the patient-level mean from one independent case (n = 2 matched patient cases); circles and triangles distinguish the individual cases, and lines connect matched non-malignant and malignant tissues from the same patient. Multiple tissue sections, acquired images, and cellular regions of interest contributed to each patient-level value.

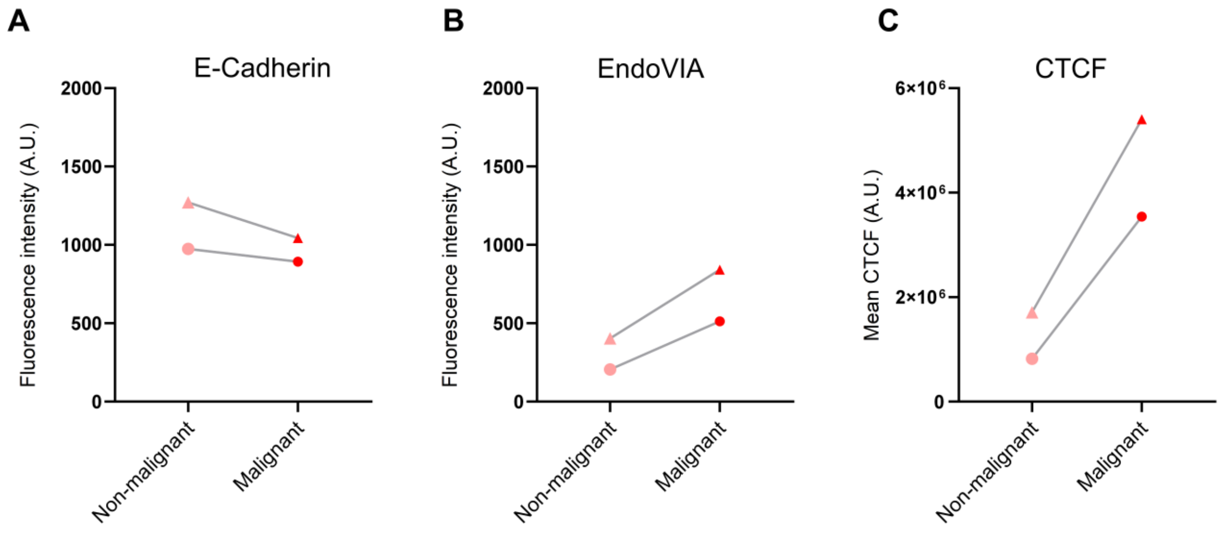


**Figure S13.** Analysis of E-cadherin and EndoVIA 2.0 fluorescence in breast FFPE tissues. (A) Mean E-cadherin fluorescence intensity in matched non-malignant and malignant breast tissues. (B) Mean EndoVIA fluorescence intensity in the same matched tissues. (C) Mean CTCF of EndoVIA in matched non-malignant and malignant tissues. Each point represents the patient-level mean from one independent case (n = 2 matched patient cases); circles and triangles distinguish the individual cases, and lines connect matched non-malignant and malignant tissues from the same patient. Multiple tissue sections, acquired images, and cellular regions of interest contributed to each patient-level value.

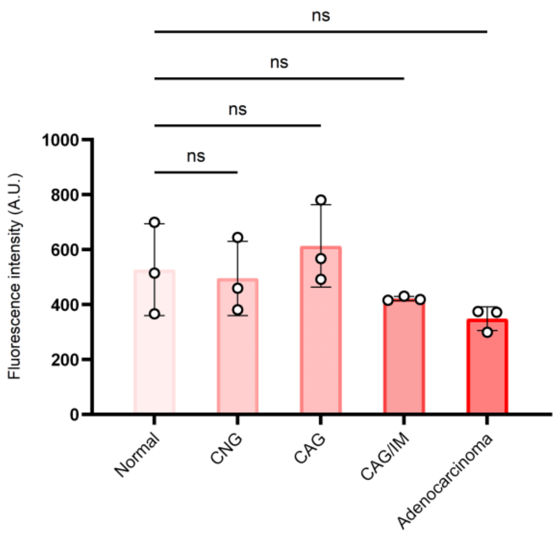


**Figure S14.** Analysis of E-cadherin fluorescence across the gastric neoplastic cascade. Mean E-cadherin fluorescence intensity in normal gastric epithelium, chronic non-atrophic gastritis (CNG), chronic atrophic gastritis (CAG), CAG with intestinal metaplasia (CAG/IM), and gastric adenocarcinoma tissues. Each point represents the patient-level mean from one independent case (n = 3 patients / group; 15 patients total). Multiple tissue sections, acquired images, and cellular regions of interest contributed to each patient level value. Data are shown as mean ± s.d. Statistical analysis was performed using Welch's one-way ANOVA. Adjusted *P* values are reported for post-hoc comparisons; ns, not significant; **P* < 0.05, ***P* < 0.01.

### Supplementary Tables

| Method | Solution | Composition |
|---|---|---|
| EndoVIA 2.0 in Cells | Permeabilization buffer | 0.1% Triton X-100, 1X PBS |
| | Blocking buffer | 3% BSA, 0.1% Triton X-100, 0.1% Tween-20, 1X PBS |
| | Calcium-containing blocking buffer | 1 mM $CaCl_2$, 3% BSA, 1X PBS |
| | Calcium-containing wash buffer | 1 mM $CaCl_2$, 1X TBS |
| EndoVIA 2.0 in Gastroids | Blocking buffer | 3% BSA, 1% Tween-20, 1% Triton X-100, 1X PBS |
| | Calcium-containing blocking buffer | 1 mM $CaCl_2$, 3% BSA, 1X PBS |
| | Calcium-containing wash buffer | 1 mM $CaCl_2$, 1X TBS |
| EndoVIA 2.0 in Fresh Frozen Tissues | 1X TBS | 20 mM Tris, 150 mM NaCl, pH 7.6 |
| | Blocking buffer | 3% BSA, 0.1% Tween-20, 1X TBS |
| | Calcium-containing blocking buffer | 3% BSA, 1 mM $CaCl_2$, 1X TBS |
| | Calcium-containing wash buffer | 1 mM $CaCl_2$, 1X TBS |
| EndoVIA 2.0 in FFPE Tissues | 1X TBS | 20 mM Tris, 150 mM NaCl, pH 7.6 |
| | Sodium citrate buffer | 10 mM sodium citrate, 0.05% Tween-20, pH 6.0 |
| dSTORM Super Resolution Setup and Imaging | dSTORM imaging buffer | 0.56 mg/mL Glucose Oxidase, 34 $\mu g \cdot mL^{-1}$ Catalase, 50 mM Tris (pH 8.00), 10 mM NaCl, 10% glucose (w/v), 143 mM β-mercaptoethanol |

**Table S1.** Buffer compositions used for EndoVIA 2.0 staining and dSTORM imaging.

| Reagent | Supplier | Catalog # | Working Dilution / Concentration |
|---|---|---|---|
| Anti-MBP (mouse) | New England Biolabs<br>Ipswich, MA, USA | E8032S | *Cells* – 1:1000<br>*Tissues* – 1:1000, 1:500 |
| Anti-MBP (rabbit) | Abcam<br>Cambridge,<br>Cambridgeshire, UK | ab9084 | *Gastroids* – 1:1000, 1:500<br>*Tissues* – 1:500 |
| Anti-mCherry (goat) | Invitrogen<br>Waltham, MA, USA | PA5-143590 | *Tissues* – 1:1000 |
| Wheat germ agglutin Alexa Fluor 488 | Thermo Fisher Scientific<br>Waltham, MA, USA | W11261 | *Tissues* – 5 μg/mL |
| Anti-E-Cadherin (rabbit) | Cell Signaling Technology<br>Danvers, MA, USA | 3195S | *Tissues* – 1:400 |
| Anti-β-actin (mouse) | Invitrogen<br>Waltham, MA, USA | MA1-140 | *Tissues* – 1:1000 |
| Anti-Histone H4 (rabbit) | Cell Signaling Technology<br>Danvers, MA, USA | D2X4V | *Tissues* – 1:200 |
| GAM Alexa Fluor 488 | Invitrogen<br>Waltham, MA, USA | A-11001 | *Tissues* – 1:1000 |
| DAG Alexa Fluor 555 | Invitrogen<br>Waltham, MA, USA | A-21432 | *Tissues* – 1:1000 |
| DAR Alexa Fluor 555 | Invitrogen<br>Waltham, MA, USA | A-31572 | *Tissues* – 1:1000 |
| GAM Alexa Fluor 647 | Invitrogen<br>Waltham, MA, USA | A-21235 | *Cells* – 1:1000<br>*Tissues* – 1:1000 |
| GAR Alexa Fluor 647 | Invitrogen<br>Waltham, MA, USA | A-21244 | *Gastroids* – 1:1000<br>*Tissues* – 1:1000 |
| DAR Alexa Fluor 647 | Invitrogen<br>Waltham, MA, USA | A-31573 | *Tissues* – 1:1000 |
| Hoechst nuclear dye 33342 | Thermo Fisher Scientific<br>Waltham, MA, USA | 62249 | 1 μg/mL |
| Anti-fade mounting medium | Thermo Fisher Scientific<br>Waltham, MA, USA | P36980 | - |
| Poly-D-lysine | Thermo Fisher Scientific<br>Waltham, MA, USA | A3890401 | 50 μg/mL |
| HEK293T cells | ATCC<br>Manassas, VA, USA | CRL-3216 | - |
| G-402 cells | ATCC<br>Manassas, VA, USA | CRL-1440 | - |
| pmGFP-ADAR1-p110 plasmid | Addgene<br>Watertown, MA, USA | 117928 | - |

**Table S2.** Antibodies, dyes, and key reagents used in Methods.
